# An integrative transcriptional logic model of hepatic insulin resistance

**DOI:** 10.1101/2021.03.15.435438

**Authors:** Takumi Kitamoto, Taiyi Kuo, Atsushi Okabe, Atsushi Kaneda, Domenico Accili

**Author notes:** Corresponding Author: Takumi Kitamoto, MD, PhD.

## Abstract

Abnormalities of lipid/lipoprotein and glucose metabolism are hallmarks of hepatic insulin resistance in type 2 diabetes. The former antedate the latter, but the latter become progressively refractory to treatment and contribute to therapeutic failures. It’s unclear whether the two processes share a common pathogenesis and what underlies their progressive nature. In this study, we investigated the hypothesis that genes in the lipid/lipoprotein pathway and those in the glucose metabolic pathway are governed by different transcriptional logics that affect their response to physiologic (fasting/refeeding) as well as pathophysiologic cues (insulin resistance and hyperglycemia). To this end, we obtained genomic and transcriptomic maps of the key insulin-regulated transcription factor, FoxO1, and integrated them with those of CREB, PPAR*α*, and glucocorticoid receptor. We found an enrichment of glucose metabolic genes among those regulated by intergenic and promoter enhancers in a fasting-dependent manner, while lipid genes were enriched among fasting-dependent intron enhancers and fasting-independent enhancer-less introns. Glucose genes also showed a remarkable transcriptional resiliency, i.e., an enrichment of active marks at shared PPARα/FoxO1 regulatory elements when FoxO1 was inactivated. Surprisingly, the main features associated with insulin resistance and hyperglycemia were a “spreading” of FoxO1 binding to enhancers, and the emergence of target sites unique to this condition. We surmise that this unusual pattern correlates with the progressively intractable nature of hepatic insulin resistance. This transcriptional logic provides an integrated model to interpret the combined lipid and glucose abnormalities of type 2 diabetes.

**Significance Statement:** The liver is a source of excess lipid, atherogenic lipoproteins, and glucose in patients with type 2 diabetes. These factors predispose to micro- and macrovascular complications. The underlying pathophysiology is not well understood, and mechanistic insight into it may provide better tools to prevent, treat, and reverse the disease. Here we propose an alternative explanation for this pathophysiologic conundrum by illustrating a transcriptional “logic” underlying the regulation of different classes of genes. These findings can be interpreted to provide an integrated stepwise model for the coexistence of lipid and glucose abnormalities in hepatic insulin resistance.

**Highlights:**

- Foxo1 regulates liver metabolism through active enhancers, and hepatocyte maintenance through core promoters
- Foxo1 regulates glucose genes through fasting-dependent intergenic enhancers
- Bipartite intron regulation of lipid genes is partly fasting-independent
- Ppar*α* contributes to the transcriptional resiliency of Foxo1 metabolic targets
- Insulin resistance causes de novo recruitment of Foxo1 to active enhancers
- A stepwise model of insulin resistance

## INTRODUCTION

An impairment of the physiologic response to insulin, or insulin resistance, remains the central cause of type 2 diabetes together with declining insulin secretory capacity, and its principal unmet treatment need (1). The pleiotropic nature of insulin resistance poses a therapeutic challenge by having different effects on different organs, and different biological consequences within the same cell type, not to mention evidence of genetic heterogeneity (2). Nowhere is this challenge more apparent than at the liver, a central organ in the pathogenesis of two key abnormalities in diabetes: increased production of atherogenic lipoproteins that increase the diabetic’s susceptibility to heart disease (1); and increased glucose production, predisposing to microvascular complications (3). In addition, the progressive nature of the latter defect (4), together with declining *β*-cell function (5), likely underlies the therapeutic failure of antidiabetic drugs (6). Among drugs directly targeting hepatic glucose production, the diabetic pharmacopeia remains woefully limited to metformin (7).

Understanding whether the two central defects of hepatic insulin resistance harken back to a shared mechanism, or arise independently, has obvious implications for the discovery of new treatments (8). A useful conceptualization that has gained some consensus separates insulin signaling into FoxO1-dependent and Srebp1c-dependent branches, the former emanating from activation of Akt and allied kinases to regulate glucose metabolism, and the latter being relayed through mTOR to supervise lipid synthetic and turnover pathways (2). However, while the case for FoxO1 regulation of specific genes is strong, its genome-wide regulatory function in the broader context of the nutrient response has only been marginally addressed (9, 10). Therefore, the extent to which the lipid and glucose metabolic branches of insulin signaling share a common regulatory network remains unknown. Moreover, transcriptional networks are integrated, redundant units with overlapping functions. During fasting, as glucagon, catecholamine, and FFA levels rise, a host of factors is activated to modulate glucose and lipid mobilization. Besides FoxO, they include CREB, PPARs, CEBPs, and nuclear receptors (11). To address these questions, we undertook to generate a liver FoxO1 cistrome in different physiologic and pathophysiologic states and compare it with the CREB, PPAR*α*, and glucocorticoid receptor cistromes. By leveraging a new mouse model developed for genome-wide interrogation of FoxO1 function (12), we discovered a FoxO1 transcriptional logic that provides insight into hepatic insulin action and resistance.

## RESULTS

### *In vivo* features of hepatic FoxO1 translocation

There is a dearth of primary data on the kinetics of hepatic FoxO1 localization in response to hormones and nutrients in the living organism. To optimize conditions for genome-wide ChIP-seq, we performed immunohistochemistry in wild-type mice to determine the time- and dose-dependence of FoxO1 nucleocytoplasmic translocation in response to insulin. Insulin injection into the inferior vena cava triggered rapid FoxO1 translocation that reached a plateau by 15 minutes (Fig. S1a), with an ED50 of 0.02U/kg (plasma level 0.4 ng/mL) (Fig. S1b). In contrast, HNF4A remained nuclear throughout (Fig. S1a). Thus, FoxO1 translocation is rapid and sensitive to physiological levels of insulin.

Next, we investigated translocation in response to fasting and refeeding. Following a physiologic 4-hr fast, 1-hr refeeding induced complete FoxO1 translocation (Fig. 1a). In contrast, a prolonged, 16-hr fast resulted in decreased FoxO1 immunoreactivity. Subsequent refeeding for up to 4 hr failed to translocate residual FoxO1 to the cytoplasm, while FoxO1 immunoreactivity increased and HNF4A immunoreactivity decreased after 2-hr refeeding (Fig. S1c). The reduced protein levels and delayed translocation are likely secondary to FoxO1 deacetylation (13–15). FoxO1 localization correlated with plasma glucose and insulin levels, as well as liver Akt phosphorylation. Thus, rapid nuclear exclusion in the 4-hr-fast/1-hr-refeed design was associated with a modest rise of glucose and insulin levels (Fig. S1d) and increased Akt phosphorylation (Fig. S1e), whereas persistent nuclear localization in the 16-hr fast/4-hr refeed design was associated with hyperglycemia, hyperinsulinemia (Fig. S1d), and reduced Akt phosphorylation (Fig. S1e). Based on these findings, we selected the 4-hr fast and 1-hr refeed time points to assess the hepatic FoxO1 regulome.

**Figure 1.**
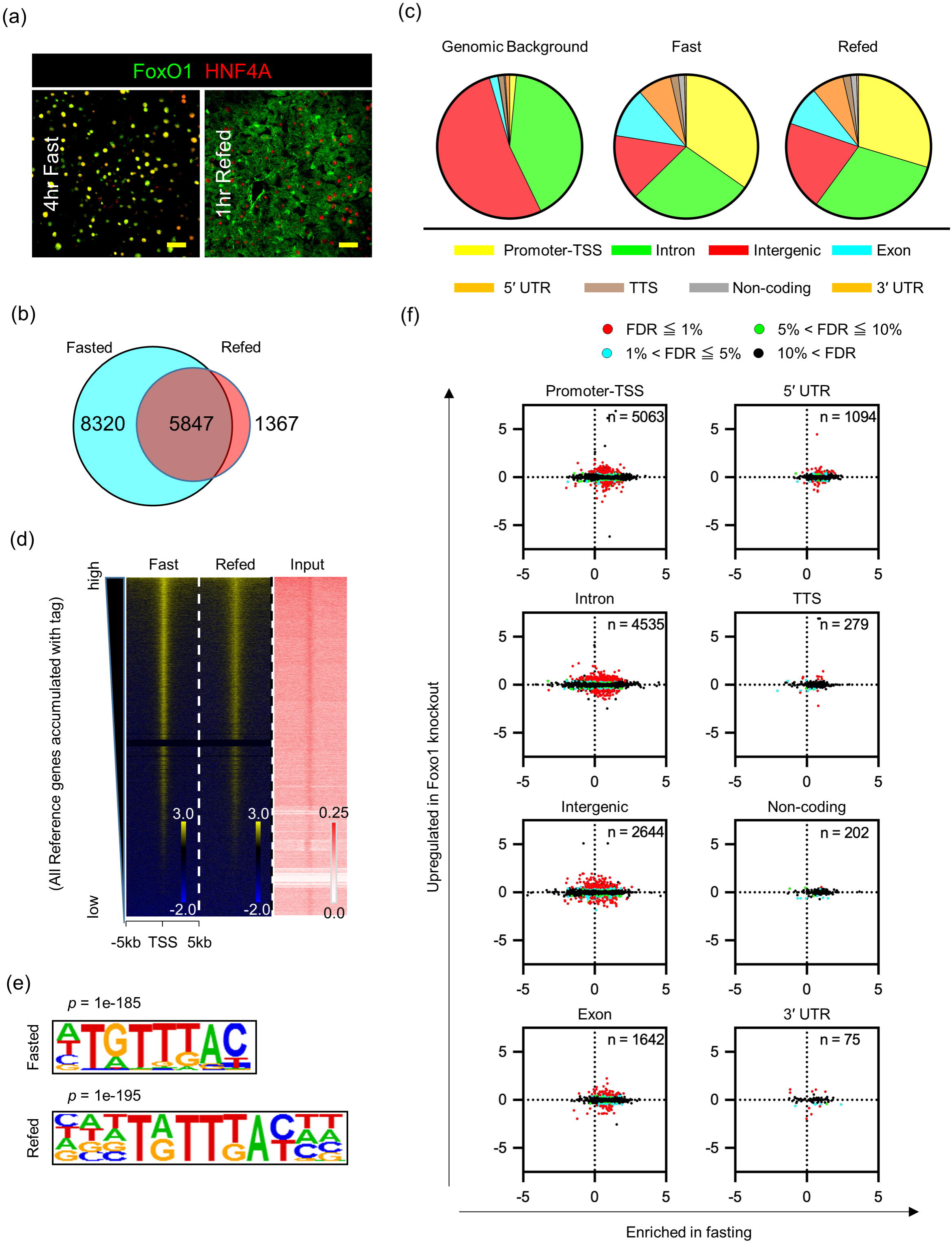
Distribution of genome-wide FoxO1 binding sites in the fast-refeed transition (a) FoxO1 and HNF4α immunohistochemistry in liver. Scale bar = 50 μm. (b) Venn diagram of the number of FoxO1 peaks in fasted or refed conditions. (c) Distribution of FoxO1 peaks relative to annotated RefSeq genes (color-coded) compared with mouse genomic background. (d) Signal intensity plots of ChIP-seq data for FoxO1 compared to input chromatin. The highest level of binding occupancy of chromatin is at the top. (e) De novo motif analysis of the FoxO1 ChIP-seq. Logos of the recovered FoxO1 motif shows position-specific probabilities for each nucleotide (*p* = 1e-185 in fast, 1e-195 in refeed). (f) Scatterplots of FoxO1 ChIP-seq peaks, expressed as log2 fold-change of FoxO1 tags between fast and refeed (horizontal axis) *vs*. log2 fold-change of mRNA levels between wild type and liver-specific FoxO1 knockout mice (vertical axis) for each genomic site. FoxO1 peaks detected in fasted or refed conditions were included in this analysis, and their number at each genomic annotation is shown inside each graph. Detailed information on peaks associated with genes whose FDR < 0.05 is in Table S1. Red= FDR < 1%; Blue= 1% ≤ FDR < 5%; Green= 5% ≤ FDR < 10%; Black= 10% ≤ FDR. See also Figure S1 and S2.

### FoxO1 regulome during fasting and refeeding

To study the genome-wide regulation of FoxO1 with fasting and refeeding, we interrogated genome occupancy by FoxO1 using an anti-GFP antibody in FoxO1-Venus knock-in mice (12) for chromatin immunoprecipitation (ChIP), to overcome the limitations of anti-FoxO1 antibodies. As reported (12), anti-FoxO1 antibodies detected the FoxO1-Venus fusion protein encoded by the modified *Foxo1* locus (Fig. S2a-b). Comparison between the two antibodies at known FoxO1 target genes (*Igfbp1, G6pc*, and *Pck1*) confirmed the specificity and superior sensitivity of the GFP antibody (Fig. S2c) (16). We next compared ChIP-qPCR and ChIP-seq using GFP antibody in FoxO1-Venus mice in the same conditions (Fig. S2d-h). Both approaches demonstrated similar decreases of FoxO1 binding to *Igfbp1*, *G6pc*, and *Pck1*, and the lack of effects on the unrelated *Fkbp5*. As the results were internally consistent, we performed further analysis with GFP antibody.

Genome-wide FoxO1 ChIP peak calling detected ∼15,000 peaks; ∼8,000 unique peaks in fasting, ∼1,000 in refeeding, and 5,000 in both conditions but to different extents (Fig. 1b). >30% of FoxO1 sites localized to promoters/transcription start sites (TSS) (Fig. 1c). Signal intensity plots demonstrated that refeeding cleared FoxO1 binding to autosomes (Fig. 1d and S3a), regardless of the distance from TSS (Fig. S3b). Known (Fig. S3c) and *de novo* motif analyses (Fig. 1e, Fig. S4) retrieved the FoxO1 motif TGTTTAC (12). This motif was found in 17% and 29.3% of FoxO1 sites in fasted and refed conditions, respectively. The same motif was found in fasted and refed conditions (Fig. S3d), and was evenly distributed between 1 and –5Kb from TSS (Fig. S3e) (17).

Next, we integrated ChIP-seq and RNA-seq data into a hepatic FoxO1 regulome. To identify FoxO1-regulated mRNAs, we induced somatic ablation of FoxO1 in liver by injecting *Foxo1^lox/lox^* mice with AAV-Cre (A-FLKO) and documented its completeness and specificity by mRNA measurements and western blotting of different tissues (Fig. S3f-g). After 3 weeks, we isolated livers from 4-hr-fasted A-FLKO and control (A-WT) mice and performed RNA-seq. We plotted the log2 difference in DNA binding (FoxO1 ChIP-seq peak number in fasted *vs*. refed animals) *vs*. the log2 difference in gene expression between A-WT and A-FLKO mice (differentially expressed genes, DEGs). Thus, the effect of genotype lies along the vertical axis, and that of fasting along the horizontal axis (Fig. 1f and Table S1).

Contingency analyses showed the strongest association between DEGs in the fasted state and FoxO1 DNA binding sites at promoters/TSS (183 of 198, or 92.4%), followed by introns (260 of 344, 75.6%), and intergenic sites (181 of 281, 64.4%), respectively (*p* < 0.0001). Since DEGs are more likely to be FoxO1 targets, these data provide initial, suggestive evidence of a FoxO1 transcriptional logic, *i.e.*, genes regulated by FoxO1 in a fasting/refeeding-dependent manner have a greater frequency of FoxO1 sites in their promoter/TSS, introns, and intergenic regions.

### FoxO1 regulates metabolic genes through active hepatic enhancers

In addition to metabolism, FoxO1 regulates cellular maintenance functions in a fasting-independent manner (18). We sought to understand the transcriptional logic of these diverging functions. We hypothesized that basic cellular functions would be regulated through core promoters, which are generally found within 1 kb from TSS and are associated with housekeeping genes and developmental TFs (19). Conversely, we surmised that metabolic genes would be regulated through tissue-specific enhancers (11, 20). To test the hypothesis, we mapped active enhancers using H3^K27ac^ and H3^K4me1^ ChIP-seq (21) in fasting and refeeding, and determined their overlap with FoxO1 sites (Fig. S5a).

Of 5,303 active enhancers co-localizing with FoxO1 sites genome-wide, 2,975 were unique to fasting, 1,022 to refeeding, and 1,306 were found in both conditions (Fig. S5b). FoxO1 enhancers localized mostly to intergenic regions and introns, and to a lesser extent to promoter/TSS (Fig. 2a). The rate of clearance in response to refeeding varied according to genomic annotation: 59.6% in intergenic regions (804/1348); 67.9% in introns, (1085/1597); and 81.5% in promoters/TSS (564/692) (*p* < 0.0001).

**Figure 2.**
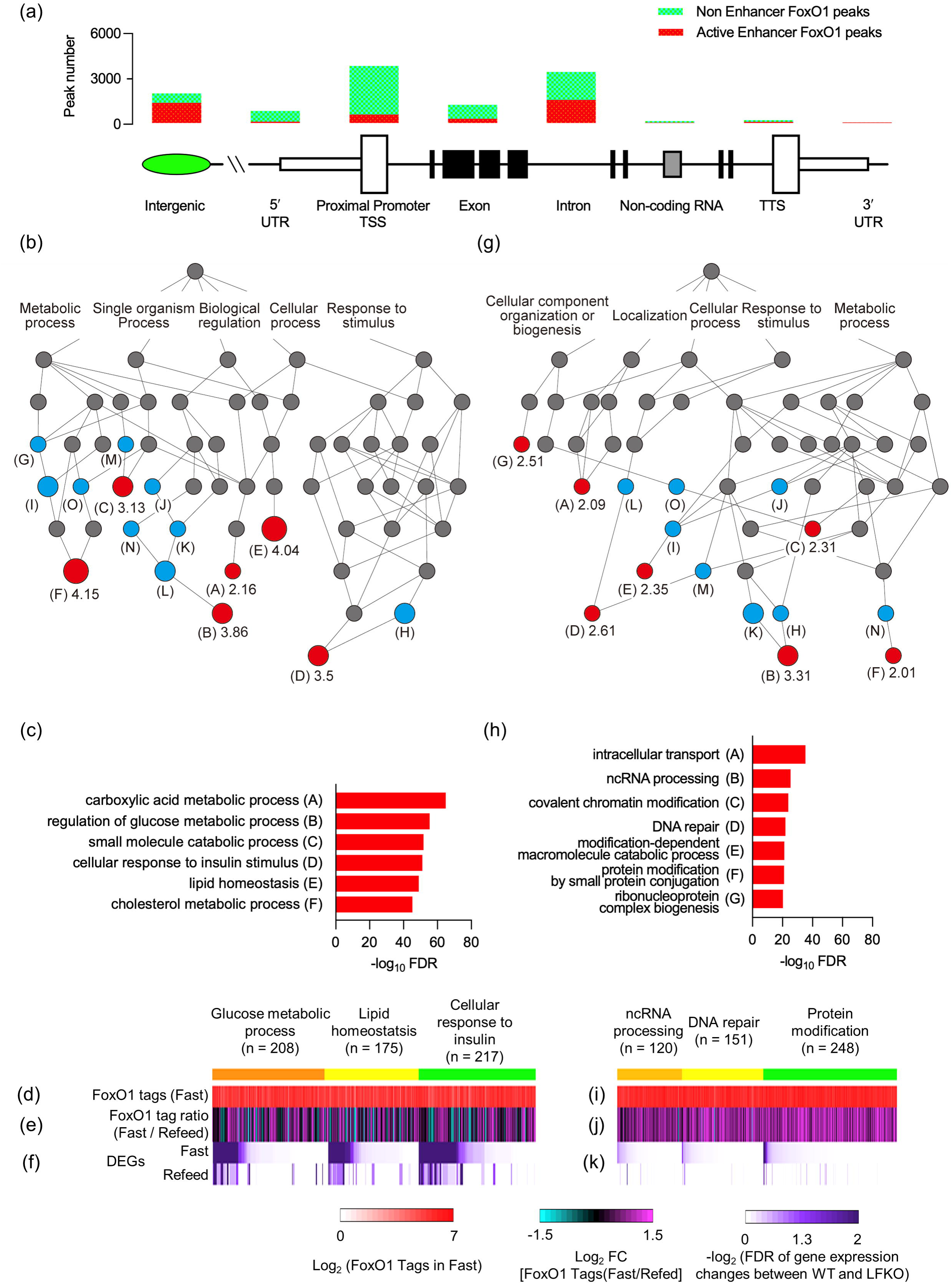
Comparison of the features of FoxO1 sites in active enhancers *vs*. non-enhancers in promoter/TSS (a) Bar diagram of FoxO1 active enhancers (red) and FoxO1 non-active enhancers (green) in each genomic location. The number of active enhancer/non-active enhancer at each genomic location is: Intergenic=1795/849, 5′ UTR= 128/966, Promoter/TSS= 760/4303, exon= 384/1258, intron= 2034/2501, non-coding= 44/158), TTS= 105/17), 3′ UTR= 53/22. (b) Directed acyclic graph derived from gene ontology analysis (GO) of biological processes associated with 5,305 FoxO1 active enhancers by GREAT GO tools. Letters correspond to the groups shown in (c) and Fig. S3c. Numbers indicate the term’s fold-enrichment. Red circles: fundamental ontologies in the hierarchy listed in (c). Blue circles: additional enriched ontologies. Gray circles: parent ontologies. (c) List of GO in (b) and their –log10 FDR. (d–f) Heatmap alignments of ChIPseq FoxO1 binding in fast (d), fast/refeed ratio (e), and FDR of gene expression changes between wild type and liver FoxO1 knockout mice (f) in GO related to glucose metabolic processes, lipid homeostasis, and cellular response to insulin genes as listed in (b, c). (g, h) Same GO analysis as in (b, c) applied to 4,303 FoxO1 sites lacking active enhancer marks in promoter/TSS. (i–k) Heatmap alignments as in (d-f) of GO related to ncRNA processing, DNA repair, and protein modification as listed in (g, h). See also Figure S3

Next, we performed ontology analyses of genes associated with FoxO1 sites in active enhancers *vs.* core promoters and visualized causal relationships among enriched terms in directed acyclic graphs (DAG) (22). FoxO1 sites in active enhancers were overwhelmingly enriched in metabolic genes, with the top three fundamental ontologies being glucose metabolism, lipid homeostasis, and insulin response (FDR 10^-40^ to ^-70^) (Fig. 2b, c, S5c). These gene ontologies showed a strong correlation between the fasting/refeeding ratio of FoxO1 DNA binding (Fig. 2d, e and S5d) (*b* = 0.09, *p* < 0.0001), and changes to mRNA expression following FoxO1 ablation, especially in fasting conditions (Fig. 2f). In contrast, enhancer-less FoxO1 sites in promoter/TSS included gene ontologies related to intracellular transport, DNA repair, ncRNA processing, and protein modification by small protein conjugation (Fig. 2g, h, S5e) (FDR 10^-20^ to^-40^). These sites showed a lesser correlation between the fasting/refeeding ratio of FoxO1 binding (Fig. 2i-j and S5f) (*b* = 0.29, *p* < 0.0001). More importantly, mRNAs encoded by genes lacking active enhancers were largely unaffected by FoxO1 ablation (Fig. 2k). The active enhancer marker, H3^K27ac^, was unaffected by fasting and refeeding (*b* = 0.91, *p* < 0.0001) (Fig. S5g).

These results indicate that the cell maintenance and metabolic functions of FoxO1 are ruled by distinct transcriptional logics: the former are governed by core promoters in a fasting/refeeding-independent manner, whereas the latter are governed by active enhancers and show a strong dependence on nutritional status (18).

### Enrichment of FoxO1 sites in introns of triglyceride and cholesterol genes

The second most common genomic annotation of FoxO1 binding sites mapped to introns (Fig. 1c). The corresponding genes showed changes to their mRNAs following FoxO1 ablation (Fig. 1f and Table S1). To understand the functional correlates of FoxO1 binding to introns, we compared expected and actual distribution of FoxO1 sites across the genome for different gene ontology groups. Interestingly, triglyceride metabolism genes showed a skewed distribution, with FoxO1 binding sites occurring at two-to three-fold the expected frequency at two locations: 5 to 50kb and –50 to –5kb from TSS (proximal introns and distal promoters), and 30 to 50% of the expected frequency at 0 to 5kb and 50 to 500kb from TSS (Fig. 3a). In contrast, glucose metabolism genes showed an enrichment 50 to 500kb from TSS, followed by the 5 to 50kb regions (Fig. 3b). Statistical analyses of annotation distribution demonstrated that triglyceride metabolism genes were significantly enriched in introns, while glucose metabolism genes were enriched in intergenic and promoter/TSS sites (*p* = 0.03) (Fig. 3c).

**Figure 3.**
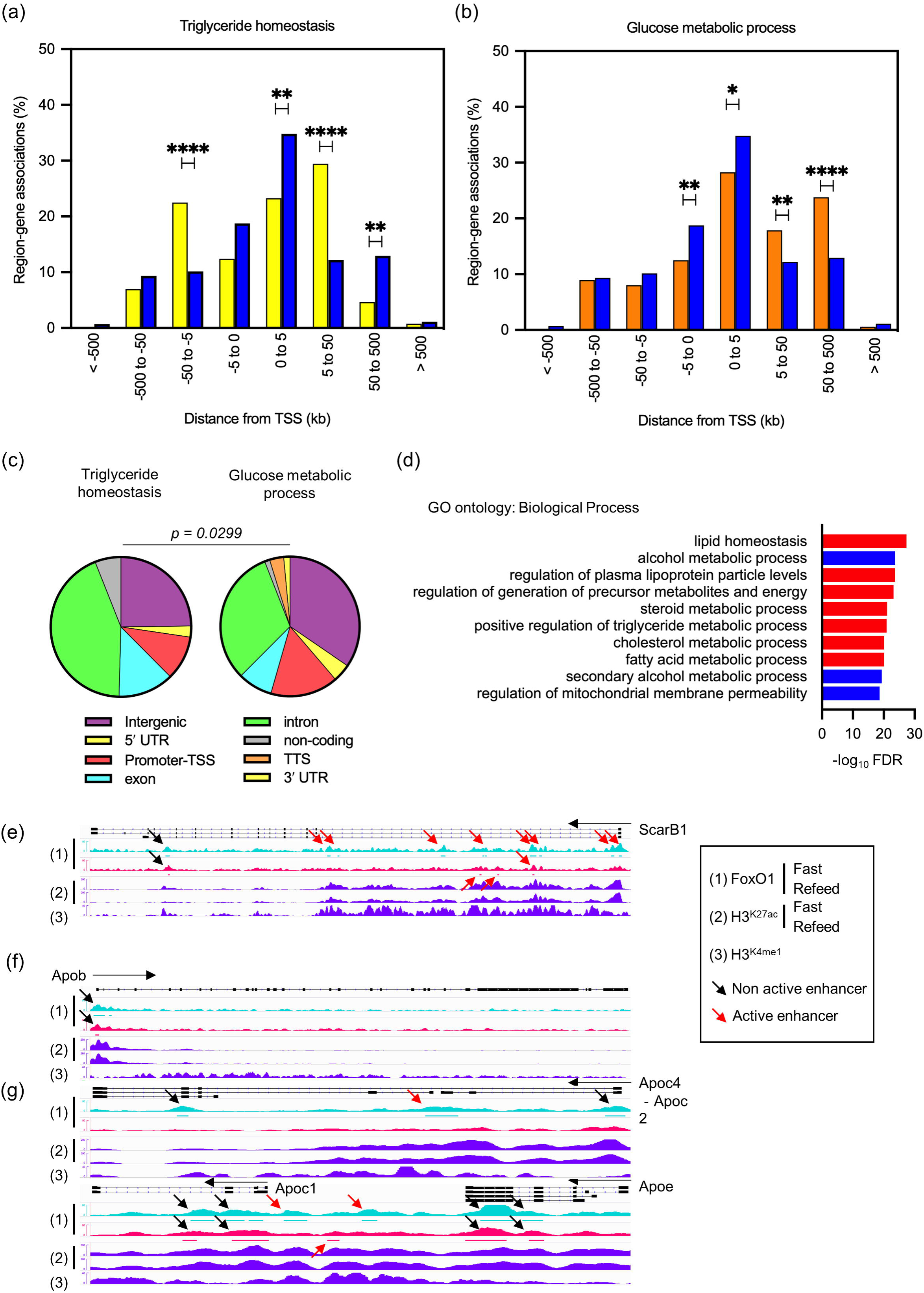
Different FoxO1 binding logic between triglyceride and glucose metabolism genes (a, b) Comparison between region-gene associations of triglyceride homeostasis (yellow bar) (a), or glucose metabolic process (orange bar), with set-wide FoxO1 binding sites (blue bar) as detected by FoxO1 ChIP-seq in fasted or refed conditions, binned by orientation and distance from TSS. *= *p* < 0.05; **= *p* < 0.01; ****= *p* < 0.0001 by chi-square test. (c) Distribution of FoxO1 binding sites associated with triglyceride homeostasis or glucose metabolic process genes according to genomic annotation as in Fig. 1c. *p=* <0.03 by contingency analysis. (d) GO analysis of biological processes associated with 4,535 FoxO1 binding sites in introns using GREAT GO tools. (e–g) IGV Genome browser views of FoxO1 peaks and associated H^3K27ac^ and H3^K4me1^ histone marks at selected apolipoprotein clusters (Apob and ApoC2/C4/C1/E Apob, Apoc4-c2, Apoc1, Apoe) and ScarB1. Signals are normalized for the comparisons between fasted and refed conditions. FoxO1 signals are aligned with peak regions. Red arrows indicate active enhancers as detected by H^3K27^ac and H3^K4me1^ signals. FoxO1 peaks in introns are listed in Table S2. See also Figure S4.

The ontology groups of intron-enriched genes included a nearly exclusive assortment of lipid, lipoproteins, and cholesterol genes (Fig. 3d). Nearly half of intron sites were associated with active enhancers (Fig. 2a). Next, we analyzed the FoxO1 regulome by intron enhancer status. Linear regression analysis of FoxO1 tags in the fasted vs. refed state demonstrated that introns marked by active enhancers showed a three-fold lower coefficient of variation than enhancer-less introns (*b* = 0.19 *vs*. 0.06) (Fig. S6a–b), and were more frequently associated with variations of the encoded mRNAs in A-FLKO. For example, ScarB1 (23) (Fig. 3e), Angptl4, and Angptl8 (24) (Table S2) showed fasting-induced FoxO1 binding to active intron enhancers, as well as altered mRNA levels upon FoxO1 ablation. In contrast, the *ApoB*, *ApoA1/C3/A4/A5* and *C2/C4/C1/E* clusters (the latter syntenic with the human *APOCII* enhancer) (25) showed fasting-independent FoxO1 binding to enhancer-less introns, and preserved mRNA expression following FoxO1 ablation (Fig. 3f-g, Table S2).

The transcriptional logic of the FoxO1 regulome emerging from the preceding analyses suggests that a majority of glucose metabolism genes are governed by an intergenic/proximal promoter/TSS active enhancer-logic in a fasting-inducible manner, whereas a majority of triglyceride, lipoprotein, and cholesterol genes are ruled by a bipartite intron-logic: fasting-dependent active intron enhancers and fasting-independent enhancer-less introns.

We hypothesized that this differential logic underlies hepatic insulin resistance. We tested the hypothesis using three different conditions: (*i*) resilience analysis to determine whether these two regulatory modalities affect the ability of these genes to undergo compensatory changes in response to variations in FoxO1 function, as a surrogate measure of insulin action (Fig. S1); (*ii*) comparative genomic analyses with other fasting-induced TFs to identify functional partners and redundancies; and (*iii*) genome-wide FoxO1 ChIP-seq in insulin-resistant/hyperglycemic mice.

### Transcriptional resiliency of glucose metabolic genes

First, we sought to determine whether different modalities of FoxO1 regulation (intergenic and promoter/TSS *vs*. intron) were associated with differential compensation by other TFs that may affect the pathophysiology of insulin resistance. To this end, we compared gene expression differences between constitutive *vs*. adult-onset somatic ablation of FoxO1 in liver (26–28) and correlated these differences with ChIP-seq data.

We generated *Alb-Cre:FoxO1^fl/fl^* mice to induce constitutive hepatic FoxO1 ablation (C-FLKO) and compared gene expression differences between adult-onset (A-FLKO, described in Fig. 1) and constitutive (C-FLKO) knockouts according to nutritional state (fast *vs*. refeed), genotype (WT *vs*. FoxO1 ablation), and timing of ablation (A-FLKO *vs*. C-FLKO) using RNA-seq (Fig. S7). t-SNE plots showed large differences in fasted *vs*. refed gene expression patterns between A-FLKO and their matched controls (A-WT). In contrast, the differences between C-FLKO and C-WT were considerably blunted (Fig. 4a). We calculated fold-change and average gene expression in each WT/knockout pair to draw MA-plots of log-intensity ratios (M-values) *vs*. averages (A-values). The number of differentially regulated genes in fasted C-FLKO mice decreased by 60% compared to A-FLKO (227 *vs.* 585), whereas it was similar in refed conditions (301 *vs*. 243) (Fig. 4b–e, Table S3). Thus, a first conclusion is that chronic compensatory changes partially mask the effect of FoxO1 ablation on the fasting response.

**Figure 4.**
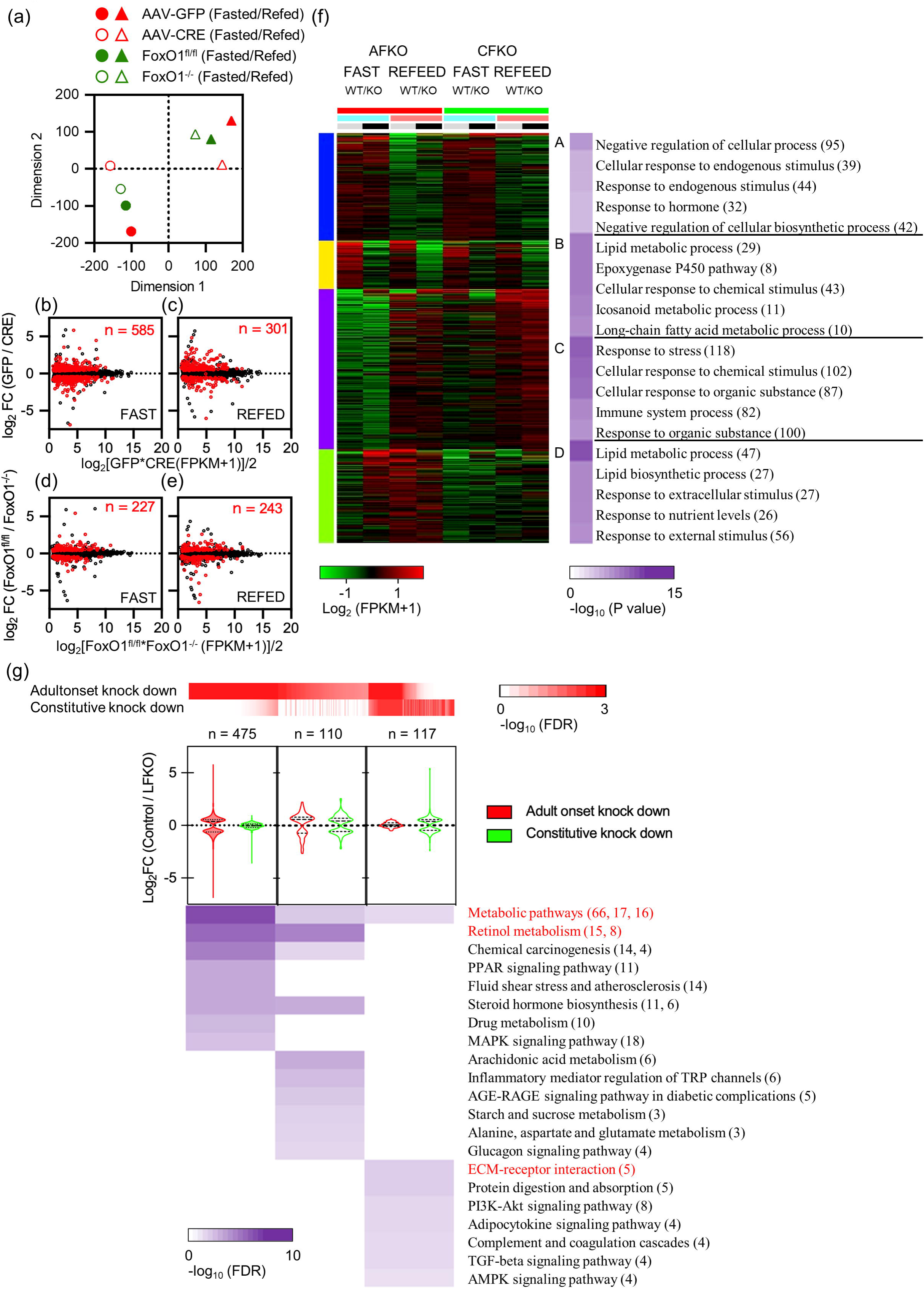
Resilience analysis of FoxO1-regulated genes (a) t-SNE plot of RNA-seq data (n= 8). Circles indicate fasted and triangles refed animals. Filled red symbols: AAV-GFP-injected animals (A-WT in the text); empty symbols with red border: AAV-CRE-injected animals (A-FLKO in the text); green filled symbols: *Foxo1^loxp/loxp^* (C-WT in the text); empty symbols with green border: Alb-Cre/*Foxo1^flox/flox^* (C-FLKO in the text). (b-e) MA-scatterplots of average expression levels vs. log2 fold-change induced by FoxO1 ablation in tag count within exons of Ensemble gene bodies in fasted (b) or refed (c) A-FLKO, and fasted (d) or refed (e) C-FLKO. Red dots represent differentially expressed genes (DEGs) (FDR ≤ 0.05). The number of DEGs is indicated in each box. (f) Enrichment analysis of k-Means clusters with molecular pathways underlying each category with top 1,000 variable genes among all samples used in (a) by iDEP tools. (g) GO analysis of DEGs in fasted conditions, shown in (b) and (d), by Shiny GO tools. Red heatmap shows FDR of genes in A-FLKO or C-FLKO. Violin plots show log2 fold-change of gene expression between control and A-FLKO (red) or C-FLKO (green) for DEGs. Number of DEGs is indicated at the top. Purple heatmap shows FDR of each ontology described next to it. Red-colored ontologies indicate the top enriched term in each category. The number of genes in each ontology is shown in parenthesis in (f, g). DEGs are listed in Table S3. See also Figure S5.

Next, we determined the ontologies of genes undergoing compensatory changes as a function of nutritional status (fast *vs*. refeed), genotype (knockout *vs*. WT), and timing-of-ablation (A-FLKO *vs*. C-FLKO) (Fig. 4f). We identified four ontology groups (A-D). Group A was comprised of genes induced by fasting, and group C of genes induced by refeeding, neither of which was affected by FoxO1 ablation in either A-FLKO or C-FLKO mice. These groups included cellular, immune, chemical, and stress response genes. In contrast, group B was comprised of genes affected by genotype (A-FLKO *vs*. A-WT and C-FLKO *vs*. C-WT), regardless of the timing of ablation. This group included primarily lipid and fatty acid metabolism genes whose expression decreased with fasting in FoxO1 knockouts. Group D was enriched in genes regulated by fasting, genotype, and timing of ablation. These genes were induced by fasting in WT, but not in A-FLKO mice. However, the differences between WT and A-FLKO were virtually lost in C-FLKO mice. This group included metabolic pathways, retinol and PPAR signaling, and steroid function genes (Fig. 4g). In contrast, only a small number of genes, primarily linked to extracellular matrix-receptor interaction and protein digestion and absorption, were uniquely affected following constitutive ablation.

We examined group D at a more granular level to identify genes in which the effect of FoxO1 ablation became less marked in C-FLKO (i.e., lower fold-change and higher FDR value between control and KO mice in C-FLKO than those in A-FLKO). These genes involved classical FoxO1 targets regulating insulin signaling (*Irs2*), gluconeogenesis (*G6pc*, *Pck1*, *Ppargc1a*), glycolysis (*Gck, Pfkfb1* and *3, Ldhd*), ketogenesis (*Hmgcs1*), and glucose/fatty acid partitioning (*Pdk4*) (Table S3). Other genes undergoing compensation included 17 members of the *Cyp2* family and 6 members of the *Cyp4* family of drug metabolizing enzymes, *Angptl8, Fgf21, Gdf15, Klf15, Slc13a5* (encoding INDY)*, Enho* (encoding Adropin)*, Fmo3,* and *Asns*.

Among genes involved in fatty acid synthesis or oxidation, apolipoproteins, and cholesterol trafficking, only *Vldlr* and *Lpin1* showed >50% compensation. Thus, FoxO1-regulated glucose metabolism genes as well as several metabolically important genes undergo a compensatory response following constitutive FoxO1 ablation, whereas the majority of lipid metabolism genes don’t. We termed this finding transcriptional resiliency.

### A FoxO1/PPAR*α* signature of fasting-inducible enhancers

Transcriptional regulation of the fasting response involves several TFs, including CREB, GR, and PPAR*α* (11). To understand the integration of these networks with FoxO1 and their potential role in the transcriptional resiliency observed after FoxO1 ablation, we compared the present dataset with published genome-wide ChIP-seq of these three factors (29, 30). Analyses of peak distribution demonstrated that CREB peaks are enriched at promoters, while GR and PPAR*α* are enriched in introns and intergenic regions (Fig. 5a). When overlaid with FoxO1 sites, we found that co-localization of FoxO1/PPAR*α* (Fig. 5b) prevailed at active intergenic and intron enhancers, where approximately half of FoxO1 sites are shared with PPAR*α* (Fig. 5c, e). In contrast, trinomial combinations FoxO1/CREB/PPAR*α* prevailed at enhancer-less promoters (Fig. 5d, e). At active enhancer sites, 11.2% of unique FoxO1 sites were associated with changes in gene expression following FoxO1 ablation, whereas only 5.4% of shared sites (FoxO1 and CREB or PPAR*α*) did (p < 0.0001, Table S4). This difference was not seen in non-active enhancer sites (6.09% vs. 5.93%, respectively) (p = NS, Table S4). Gene ontology analysis (Fig. 5f) showed that abnormal gluconeogenesis is the most significant annotation of FoxO1/PPAR*α* shared intergenic peaks (FDR = 2.22 × 10^-31^), while lipid homeostasis is the most significant in introns (FDR = 2.01 × 10^-21^).

**Figure 5.**
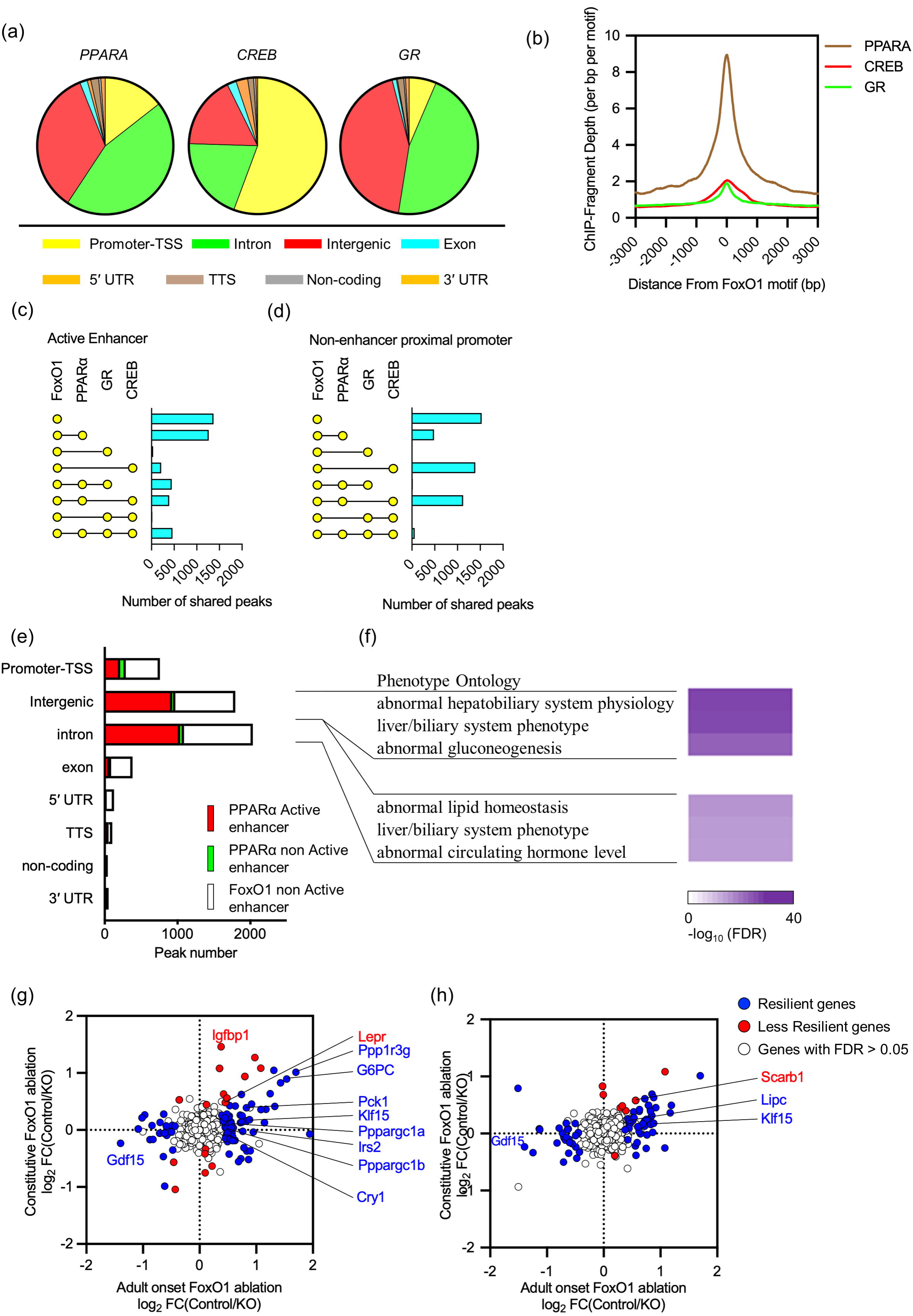
Comparative analysis among fasting inducible transcriptional factors (a) Distribution of PPARα, CREB and GR binding sites in fasted conditions. (b) Peak plot mapping the overlap of the FoxO1 (Fig. 1e) and PPARα, CREB and GR peaks. (c-d) Intersection analyses of active (c), and non-active (d) FoxO1 and PPARα, CREB or GR enhancer peaks in fasting conditions. (e) Proportion of PAARα peaks with/without active enhancer marks in FoxO1 active enhancers in fasting conditions according to genomic annotation. (f) Heatmap with associated FDR of phenotype ontology terms of shared FoxO1/PPARα active enhancers (red bars) in intergenic regions and introns. (g, h) Resiliency plots of genes associated with shared FoxO1/PPARα active enhancers in intergenic regions (g) and introns (h). Plot show log2 fold-change induced by adult-onset vs. constitutive liver FoxO1 ablation. Resilient genes (FDR ≤ 0.05 in AFKO or CFKO mice, showing lower fold-change and higher FDR value in CFKO mice than AFKO mice) are indicated by blue dots, non-resilient genes (FDR ≤ 0.05 in AFKO or CFKO mice) are marked by red dots, FDR > 0.05 in both mice by white dots. DEGs are listed in Table S5.

Next, we asked whether co-regulation by FoxO1 and PPAR*α* can explain the resiliency of gene expression. We plotted each FoxO1/PPAR*α* shared peak with active enhancer marks *vs*. changes to mRNA encoded by associated genes in A-FLKO and C-FLKO (Fig. 5g-h). In both intergenic (Fig. 5g) and intron (Fig. 5h) sites, >80% of FoxO1/PPAR*α* co-regulated genes showed a compensatory response to constitutive FoxO1 ablation (75 of 92 and 68 of 76, respectively). In intergenic sites, we found notable resilient glucose metabolism genes, such as *Pck1*, *G6pc, Irs2*, *Ppargc1a and b*, *Ppp1r3g, Cry1, Gdf15* (31) and *Klf15* (32) (Fig. 5g). In introns, we found lipid genes, such as *Gdf15,* and *Lipc* (Fig. 5h, Table S5). Thus, shared FoxO1/PPAR*α* enhancers are more likely to undergo compensation when FoxO1 is inactive.

### Enhancer spreading of FoxO1 binding in insulin resistance/hyperglycemia

To evaluate the effects of insulin resistance and hyperglycemia on the FoxO1 regulome, we subjected FoxO1-Venus mice to high fat diet (HFD) or treatment with the insulin receptor antagonist, S961 (33). Both interventions impaired refeeding-induced FoxO1 translocation (Fig. 6a) and caused hyperglycemia (not shown). However, as the effects of S961 were more marked, we performed genome-wide ChIP-seq in livers of 4-hr-fasted/1-hr-refed mice treated with S961 *vs*. vehicle.

**Figure 6.**
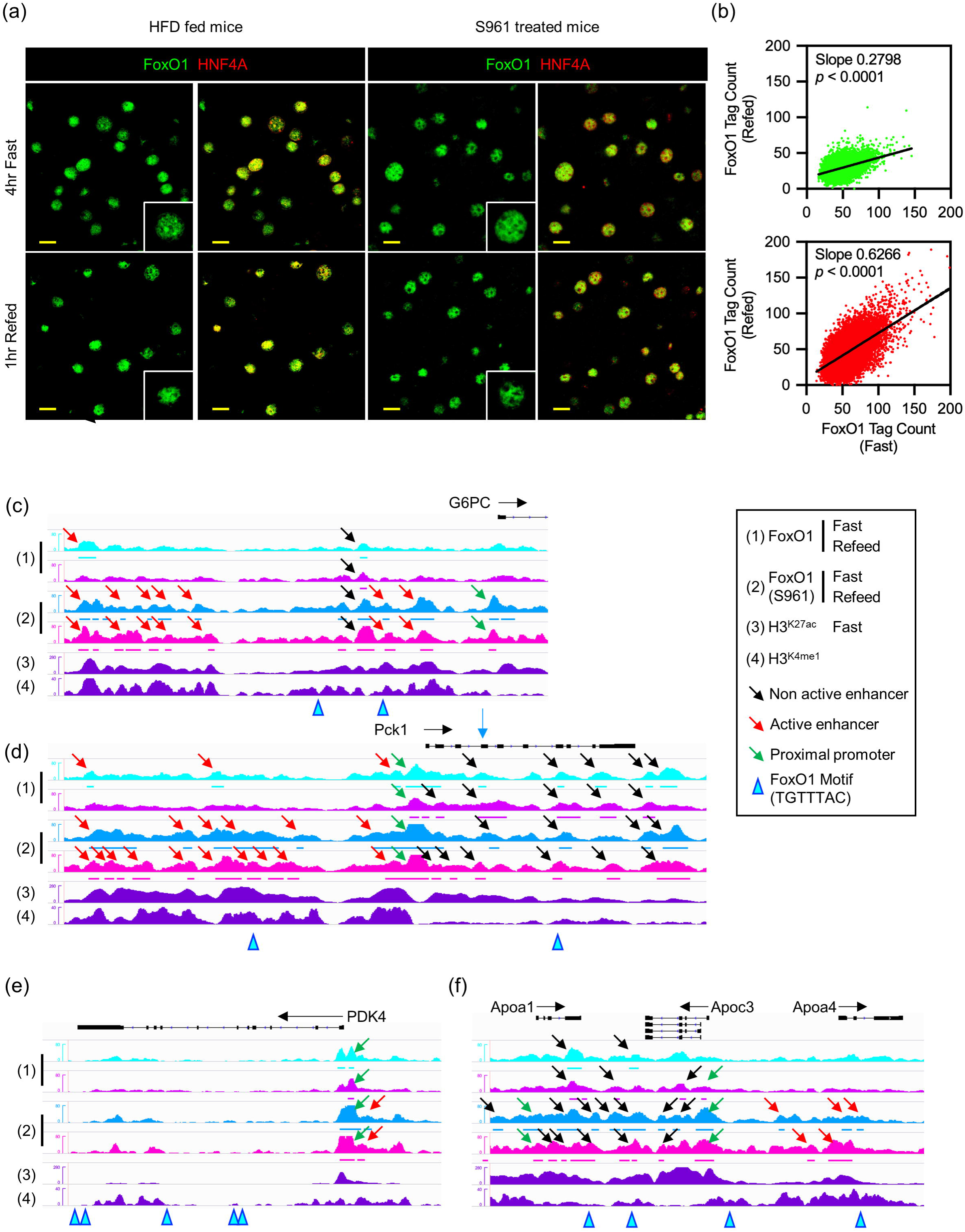
The transition of FoxO1 binding sites under insulin resistant condition (a) Immunohistochemistry of FoxO1 and HNF4α after 4hr fasting or following 1hr refeeding in high fat diet (HFD)-fed mice or insulin receptor antagonist (S961)-treated mice. Scale bar = 20 μm. (b) Scatterplots showing linear regression analysis of FoxO1 tag count between fasted and refed conditions. Green: vehicle; red: S961-treated mice. (c–f) IGV Genome browser views of FoxO1 peaks with or without S961 treatment and associated H3^K27ac^ and H3^K4me1^ marks of at *G6pc, Pdk4, Angptl4/8, and ApoA1/C3/A4*. See also Figure S6.

Regression analysis of FoxO1 tags under fasted and refed conditions showed a two-fold higher coefficient in S961-treated mice than in vehicle controls (*b* = 0.28 *vs*. 0.62, Fig. 6b), consistent with impaired translocation (Fig. 6a). We examined FoxO1 binding to representative genes of the two main transcription logics identified above, intergenic/promoter/TSS (glucose) *vs*. intron (lipid) genes. We found the emergence of novel FoxO1 binding patterns at active enhancers of both classes. Examples included intergenic/promoter enhancers of glucogenic (*G6pc, Pck1*, *Klf15*) (Fig. 6c-d, S8a) and glucose–lipid metabolic partitioning genes (*Pdk4*) (Fig. 6e), as well as intron enhancers of lipid/cholesterol genes (*ApoA1/C3/A4, Scarb1*) (Fig. 6f, S8b-c). These novel FoxO1 peaks were unaffected by fasting/refeeding, and included both FoxO1 binding motif-containing sites and sites without FoxO1 motifs. In contrast, novel FoxO1 marks at enhancer-less sites occurred less frequently. Thus, insulin resistance and hyperglycemia bring about an ectopic, dysregulated binding of FoxO1 at enhancer sites, which we term enhancer spreading.

## DISCUSSION

The present study provides transcriptional logic insight into the differential regulation of glucose and lipid metabolism in response to nutrient changes, and in insulin resistance. There are obviously non-transcriptional components to this pathophysiologic state that are partly cell-nonautonomous (34), but the present study was designed to establish genome-wide map that integrates multiple TFs, including FoxO1, with the salient pathophysiologic features of hepatic insulin action and resistance. The main conclusions are: (*i*) the transcriptional logic of FoxO1 is compatible with the bifurcating model of insulin signaling to lipid *vs*. glucose metabolism (35), whereby glucose metabolic genes are governed by intergenic and promoter/TSS enhancers, and lipid genes by a bipartite intron logic that includes fasting-dependent intron enhancers and fasting independent enhancer-less introns. (*ii*) Active enhancers of glucose metabolic genes show transcriptional resiliency, likely through shared PPARα/FoxO1 regulatory elements. (*iii*) Insulin resistance and hyperglycemia result in the spreading of FoxO1 binding to enhancers, resulting in quantitative and qualitative abnormalities of FoxO1 marks (12).

Based on these findings, we propose this model (Fig. 7): in the physiologic fasting/refeeding transition, FoxO1 is cleared more efficiently from enhancer-containing sites than from enhancer-less sites. As the former are more tightly associated with glucose genes, and the latter with lipid/lipoprotein genes, in the initial stages of insulin resistance glucose genes can still be regulated, while regulation of lipid genes is impaired. This differential sensitivity can explain why lipid/lipoprotein abnormalities chronologically precede hyperglycemia in the progression of diabetes (36). Further work will be required to functionally interrogate different classes of sites. As insulin resistance progresses, the gradual compensation of glucose vs. lipid genes in response to chronic *vs*. adult-onset FoxO1 ablation (transcriptional resiliency at intergenic and promoter/TSS enhancers) can be interpreted to suggest that glucose genes can gradually become FoxO1-independent, allowing transcription factors (likely PPARα) to induce their expression. In the clinically overt stage of the disease, as insulin resistance increases, activation of FoxO1 at ectopic (or low-affinity) enhancers leads to worsening fasting hyperglycemia, and may possibly underlie therapeutic failures. The proposed model integrates *in vivo* pathophysiological and cell biological data with genome-wide assessments to explain a clinical conundrum that has important practical implications for treatment and drug development (1). This model also addresses two criticisms leveled at the FoxO-centric view of insulin action: *(i)* that FoxO1 sensitivity to insulin makes it an unlikely candidate as a mediator of insulin resistance (37); and (*ii*) that transcription of candidate glucogenic genes alone does not fully explain increased hepatic glucose production (38). Indeed, the gamut of FoxO1 targets includes most genes involved in insulin action, and the failure to detect abnormalities in their expression following constitutive somatic ablation of FoxO1 can be explained by their resiliency.

**Figure 7.**
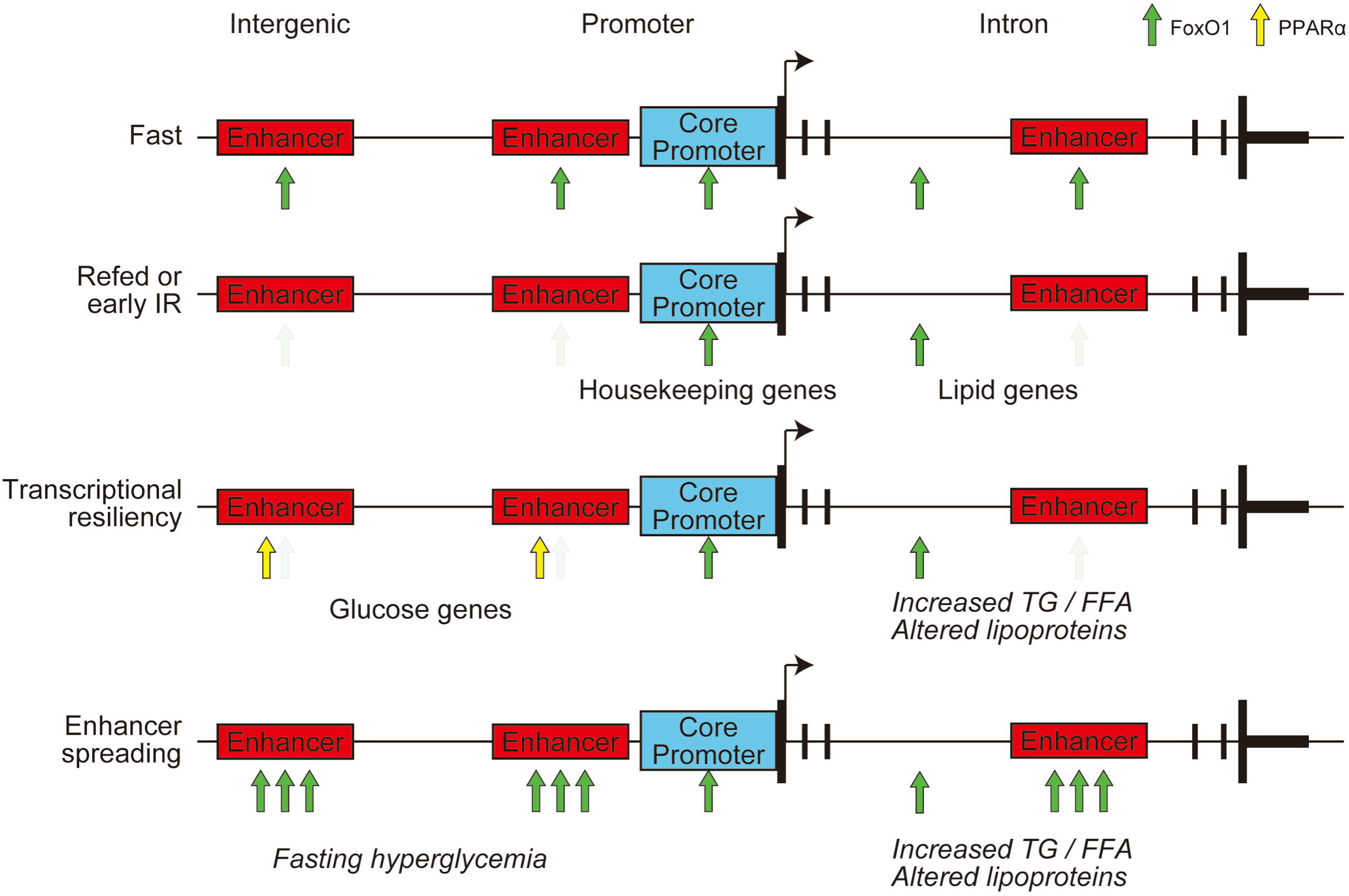
Model of FoxO1 transcriptional logic in the pathogenesis of selective insulin resistance In normal conditions, FoxO1 is cleared upon refeeding from resilient enhancers, enriched in glucose metabolism genes, but not in introns, enriched in lipid metabolism genes. With the onset of insulin resistance-induced hyperinsulinemia, FoxO1 can be cleared from resilient enhancers, but not from introns, increasing serum lipoprotein and triglyceride levels. As insulin resistance progresses, compensation by PPARα and spreading of FoxO1 binding to additional sites bolsters expression of glucose metabolic genes, inducing fasting hyperglycemia with dyslipidemia.

To demonstrate a distinctive FoxO1 transcriptional logic, we decisively leveraged the ability to examine FoxO1 targets by genome-wide ChIP-seq (12). Previous studies have been limited by the sensitivity of available FoxO1 antibodies, and have therefore detected fewer FoxO1 binding sites (9, 10, 39). There is a partial dissociation between the ChIP results, indicating that FoxO1 is still bound at several sites after refeeding, and the immunofluorescence that shows FoxO1 nuclear exclusion. However, ChIP is more sensitive than immunohistochemistry, being based on PCR amplification, and can detect lower levels of FoxO1 protein. The formation of different molecular complexes likely underlies the different modes of FoxO1 action. In this regard, we have previously shown that SIN3a is the FoxO1 corepressor at glucokinase, providing a mechanistic precedent for gene-specific targeting (8). The preferential regulation of FoxO1 by fasting/refeeding at active enhancers likely results from intrinsic and extrinsic factors, such as higher DNA accessibility at active enhancers (40), and active enhancer-promoter interactions (41) that affect assembly of pre-initiation complexes, initiation of transcription by RNA polymerase II, or transcription bursting (19).

Following FoxO1 ablation, expression of its targets can be compensated for by transcription factors acting synergistically, through its paralogue FoxO3, or reorganization of chromatin accessibility at sites where FoxO1 acts as pioneer transcription factor (42), as shown with other FoxO isoforms (27). Interestingly, genes associated with glucose metabolism (*G6pc, Pck1, Ppargc1a, Pdk4* and *Klf15*), but not those regulating general cellular responses, are selectively compensated for following FoxO1 ablation. FoxO1 peaks in these genes are cleared by refeeding, but not in insulin-resistant conditions. These genes have been shown to play a role in diabetes in studies with insulin-resistant mice (26, 43–45).

There are parallels between our findings and recent evidence that immunocyte differentiation is controlled by an enhancer- or core promoter-driven logic, with a striking partition between the two gene sets (46). The former activity is cued by the overall activity pattern of distal enhancers, while the latter is aligned with promoters. Although it is disputed whether core promoters and enhancers represent different entities or synergistically regulate transcriptional bursting, enhancers are thought to be tissue-specific, and thus more likely to confer specificity on the tissue-specific metabolic functions of FoxO1 (20).

Our comparative analysis provides evidence of cooperative and non-cooperative interactions with GR, CREB and PPAR*α*, the latter involving up to half of the FoxO1 sites in active enhancers. The extensive sharing of intergenic active enhancers of glucose genes by FoxO1 and PPAR*α* is a novel finding of this study that dovetails with the different physiologic cues regulating these two TFs. During fasting, glycogenolysis precedes gluconeogenesis and the generation of FFA substrates that activate PPAR*α* (47). Thus, we envision that FoxO1 and PPAR*α* act in a physiologic relay to ensure continuity between the early and late fast. The significant overlap between FoxO1 and PPARα may also provide an explanation for the relatively mild phenotypes of liver-specific inactivation of FoxO1 (26) and PPAR*α* (48). Functional elucidation of their interactions will be important to determine key targets in glucose metabolism and their role in diabetes pathogenesis.

## Data sharing

Further information and requests for resources and reagents should be directed to and will be fulfilled by Takumi Kitamoto.

## Data and Code Availability

The ChIP-seq and RNA-seq datasets generated during this study are available at the NCBI GEO [GSE151546]

## EXPERIMENTAL MODEL AND SUBJECT DETAILS

### Animals

Mice were housed in a climate-controlled room on a 12h light/dark cycle with lights on at 07:00 and off at 19:00, and were fed standard chow (PicoLab rodent diet20, 5053; PurinaMills). Male mice of C57BL/6J background aged 8-12 weeks were used. FoxO1-Venus mice have been described (12, 49). Briefly, To express GFP (Venus), we obtained the pCAG:myr-Venus plasmid. A 15-amino acid linker sequence was placed between the C terminus of FoxO1 and N terminus of Venus to alleviate steric hindrance. We used BAC recombineering to generate FoxO1-Venus ES cells. To generate constitutive liver-specific FoxO1 knockouts, we crossed FoxO1^lox/lox^ and Albumin-cre (50) transgenic mice. Adult onset liver-specific FoxO1 knockout mice were generated by injection of 1×10^11^ purified viral particles (AAV8.TBG.eGFP or AAV8.TBG.Cre, Penn Vector Core) per mouse via tail vein. We performed metabolic analysis or killed animals on day 21 post-injection. To assess FoxO1 localization and other liver parameters, we took organs from 4-hr-fasted (10:00 to 14:00) or 4-hr-fasted/1-hr-refed mice. For prolonged fasting experiments, we removed food overnight (18:00 to 10:00). Mice were killed 0, 1, 2, or 4 hr after refeeding. For insulin treatment, we anesthetized 16-hr-fasted mice with ketamine (100mg/kg) and xylazine (10mg/kg) i.p., followed by injection of 1U/kg insulin (NovoLog®, Novo Nordisk, Denmark) in the inferior vena cava (IVC). We collected blood and took the liver before and after insulin injection. Blood glucose was measured using (CONTOUR^®^NEXT ONE, Ascensia, USA), and insulin with a mouse-specific ELISA kit (Mercodia, USA). All animal experiments were in accordance with NIH guidelines, approved and overseen by the Columbia University Institutional Animal Care and Use Committee.

### Primary hepatocyte isolation

Primary hepatocyte isolation was performed as described (51). We anesthetized male mice with ketamine (100mg/kg) and xylazine (10mg/kg) i.p., cannulated the IVC with a 24-gauge catheter (Exel international), and infused 50 cc EGTA-based perfusion solution followed by 100 cc type I collagenase solution (Worthington Biochemicals). Following cell dissociation, we filtered cells with 100 mm mesh cell strainers, and gradient centrifugation steps to purify cell suspension. Then, we suspended hepatocytes at 5 × 10^5^ cells / mL in Medium 199 (Sigma), 10% FBS (Life Technologies), antibiotics (plating medium). After plating for 2 hr on collagen-coated plates, we exchanged plating medium for 4 hr.

## METHOD DETAILES

### Chemicals and antibodies

Ketamine was from KetaSet® and Xylazine from AnaSed®; medium 199, HBSS, EGTA, HEPES, PenStrep and Gentamycin from Life Technology; collagen type 4 from Worthington; Insulin (NovoLog®) and S961 from Novo Nordisk A/S; sodium orthovanadate from New England Bio; Bovine Serum Albumin (BSA) from Fisher Scientific. 16% paraformaldehyde (PFA) was from Electron Microscopy Sciences, and was diluted in sterile phosphate buffer solution to 4% final concentration. Anti FoxO1 (for Western Blot and immunohistochemistry, C29H4), anti panAkt (for Western Blot, 40D4) and phosphor-Akt (Ser473) (for Western Blot, D9E), normal Rabbit IgG (for chromatin immunoprecipitation, 2729) were from Cell Signaling. HNF4A (for immunohistochemistry, ab41898), GFP (for chromatin immunoprecipitation, ab 290), FoxO1 (for chromatin immunoprecipitation, ab39670) were from Abcam. H3K27ac (for chromatin immunoprecipitation, 39133) was from Active motif. Anti GFP (immunohistochemistry, A-6455) was from Invitrogen.

### Protein analysis

Livers were lysed in sonication buffer containing 20 mM HEPES pH7.5, 150 mM NaCl, 25 mM EDTA, 1% NP-40, 10% glycerol, 1 mM Na vanadate, 1 mM phenylmethylsulphonyl fluoride (PMSF), and protease and phosphatase inhibitors cocktail (Cell Signaling). We sonicated lysates for 100 sec (5×, output 70%, 20sec/20sec) and centrifuged them for 15 min at 14,000 rpm. 30 µg protein (Pierce BCA, Thermo scientific) was subjected to SDS-PAGE. We used the following antibodies: Akt (1:2,000), phosphor-Akt (Ser473) (1:2,000), β-actin (1:1,000), FoxO1 (1:1,000) (all from Cell Signaling), and GFP (1:1,000) (Abcam, ab290).

### Immunohistochemistry

We anesthetized 8-to 12-week-old mice fasted or refed for various lengths of time and perfused them with 4% PFA through the IVC. Livers were collected, fixed in 4% paraformaldehyde for 2-hr, dehydrated in 30% sucrose overnight at 4°C, embedded in OCT (Sakura, Torrance, CA), frozen to −80°C, and cut into 7-µm sections. We used primary antibodies to FoxO1 (1:100; Cell signaling technology, Boston, MA) and HNF4A (1:100; Abcam, Cambridge, MA), and secondary anti-IgG antibodies conjugated with Alexa Fluor 488 and 555 for each of the species (1:1,000; Life Technologies). Immunofluorescence was visualized by the TSA fluorescence system (PerkinElmer, Waltham, MA).

### Real-time qPCR

We lysed livers in 1 mL of TRIzol, purified RNA using RNeasy Mini Kit (Qiagen, Germantown, MD), reverse-transcribed it with qScript cDNA Synthesis Kit (QuantaBio, Beverly, MA), and performed PCR with GoTaq® qPCR Master Mix (Promega, Madison, WI). Primer sequences are available upon request. Gene expression levels were normalized to 18S using the 2-DDCt method and are presented as relative transcript levels.

### RNA-seq library constructions and data analysis

We prepared the samples from three mice for each group, and generated the libraries individually. Libraries for RNA-seq were prepared using the TruSeq Stranded mRNA Sample Prep Kit (Illumina), following the manufacturer’s protocol. Deep sequencing was carried out on the Illumina NextSeq 500 platform using the NextSeq 500/550 high-throughput kit v2.5 (Illumina) in 75-base single-end mode according to the manufacturer’s protocol. Sequenced reads from the RNA-seq experiment were aligned to mouse genome mm10 using HISAT2. Cufflinks was used for transcript assembly. Gene expression levels were expressed as fragments per kilobase of exon per million mapped sequence reads and Cuffdiff was used for statistical comparison.

### Chromatin immunoprecipitation assays and ChIP-seq library construction

The ChIP-IT® High Sensitivity kit (Active Motif, Carlsbad, CA) was used for chromatin immunoprecipitation (ChIP) following the manufacturer’s protocol. We anesthetized 8-to 12-week-old mice after 4-hr fasting followed or not by 1-hr refeeding and perfused them with 10 μM orthovanadate through the IVC. We harvested samples from left lobe of liver tissues and pooled 100mg of samples from three individual replicates for further experiments. We obtained sheared chromatin from 300 mg of liver extract using a S220 Focused-ultrasonicator (Covaris). Immunoprecipitation was performed using 4 µg of anti-GFP antibody for 10 µg of sheared chromatin. The specificity of the anti-GFP antibody was confirmed by western blotting of liver extract. ChIP-seq libraries were constructed using KAPA Hyper Prep Kit (KAPA Biosystems) according to the manufacturer’s instructions. ChIP-seq libraries were quantified by Tapestation (Agilent) and sequenced on an Illumina NEXTseq (Illumina, San Diego, CA, USA) with 75-base single-end mode.

### ChIP-qPCR

Real-time ChIP-qPCR was carried out as described above. The signal of binding events was normalized against input DNA for primer efficiency (Active Motif). Quantitative PCR primers used are listed. *G6pc* forward: GCCTCTAGCACTGTCAAGCAG and reverse: TGTGCCTTGCCCCTGTTTTATATG; *Pck1* forward: TCCACCACACACCTAGTGAGG and reverse: AGGGCAGGCCTAGCCGAGACG; *Igfbp1* forward: ATCTGGCTAGCAGCTTGCTGA and reverse: CCGTGTGCAGTGTTCAATGCT; Fkbp5 forward: TTTTGTTTTGAAGAGCACAGAA and reverse: TGTCAGCACATCGAGTTCAT.

### ChIP-seq data analysis

Reads were aligned to mouse genome mm10 using Bowtie2 software (52). The reads used in subsequent analysis passed Illumina’s purity filter, aligned with no more than 2 mismatches, and mapped uniquely to the genome. Duplicate reads were removed with Picard tools. The tags were extended at their 3′-ends to 200-bp. Technical information of sequencing depth and aligned reads is summarized in Table S6. Peak calling was performed by MACS 2.1.0 (53) with the *p*-value cutoff of 10^-7^ for narrow peaks and with the *q-*value cutoff of 10^-1^ for broad peaks against the input DNA control sample. The transcription start site (TSS) determined on mouse genome mm10 was used as measurement of the distance of each peak. HOMER software suite (54) was used to perform motif analysis, annotate peaks, such as promoter/TSS, introns, exons, intergenic, 5′ UTR, non-coding RNA, and 3′ UTR, merge files, and quantify data to compare peaks. For the detection of active enhancers, we used bedtools (55) by collecting the intersection of the peaks of TF and histone marks.

### In vivo insulin-resistant model

For high-fat diet-induced insulin resistance, animals were fed either standard or High-fat chow (Rodent Diet with 60kcal% fat, D12492i; Research diets Inc.) beginning at 8 weeks of age for 4 weeks. For S961 treatment, vehicle (normal saline) or 10nmol S961 was loaded into Alzet osmotic pumps 2001 and implanted subcutaneously on the back of mice. Mice were euthanized 3 days after implantation.

### Additional Data Sets

The following public source data were used in this work: ChIP-seq data from adult mouse liver [H3^K4me1^] (56) (GEO: GSE31039), PPARα (30) (GEO: GSE35262), GR and CREB (29) (GEO: GSE72084).

## QUANTIFICATION AND STATISTICAL ANALYSES

Values are presented as means ± SEM, and analyzed using Prism 8.2.1 (GraphPad Software, Inc.). We used unpaired Student’s *t*-test for normally distributed variables for comparisons between two groups, one-way ANOVA followed by Bonferroni post-hoc test for multiple comparisons, and Pearson’s correlation coefficient to investigate the relationship between two variables. Chi-square tests are applied for contingency analysis. We used a threshold of *p* < 0.05 to declare statistical significance.

## Acknowledgments

We thank members of the Accili and Kaneda laboratories, Dr. Utpal B. Pajvani, Dr. Rebecca A. Haeusler, Dr. Michael J Kraakman for insightful discussions of the data. Mr. Thomas Kolar and Ms. Ana Maria Flete (Columbia University) for exceptional technical support.

## References

1. E. Ferrannini, R. A. DeFronzo, Impact of glucose-lowering drugs on cardiovascular disease in type 2 diabetes. Eur Heart J 36, 2288–2296 (2015).

2. R. A. Haeusler, T. E. McGraw, D. Accili, Biochemical and cellular properties of insulin receptor signalling. Nat Rev Mol Cell Biol 19, 31–44 (2018).

3. R. N. Bergman, M. S. Iyer, Indirect Regulation of Endogenous Glucose Production by Insulin: The Single Gateway Hypothesis Revisited. Diabetes 66, 1742–1747 (2017).

4. R. A. Rizza, Pathogenesis of fasting and postprandial hyperglycemia in type 2 diabetes: implications for therapy. Diabetes 59, 2697–2707 (2010).

5. D. Accili, Insulin Action Research and the Future of Diabetes Treatment: The 2017 Banting Medal for Scientific Achievement Lecture. Diabetes 67, 1701–1709 (2018).

6. L. Monnier, C. Colette, G. J. Dunseath, D. R. Owens, The loss of postprandial glycemic control precedes stepwise deterioration of fasting with worsening diabetes. Diabetes Care 30, 263–269 (2007).

7. M. Evans, A. R. Morgan, Z. Yousef, What Next After Metformin? Thinking Beyond Glycaemia: Are SGLT2 Inhibitors the Answer? Diabetes Ther 10, 1719–1731 (2019).

8. F. Langlet et al., Selective Inhibition of FOXO1 Activator/Repressor Balance Modulates Hepatic Glucose Handling. Cell 171, 824–835 e818 (2017).

9. W. Fan et al., FoxO1 regulates Tlr4 inflammatory pathway signalling in macrophages. Embo J 29, 4223–4236 (2010).

10. D. J. Shin et al., Genome-wide analysis of FoxO1 binding in hepatic chromatin: potential involvement of FoxO1 in linking retinoid signaling to hepatic gluconeogenesis. Nucleic Acids Res 40, 11499–11509 (2012).

11. I. Goldstein, G. L. Hager, Transcriptional and Chromatin Regulation during Fasting - The Genomic Era. Trends Endocrinol Metab 26, 699–710 (2015).

12. T. Kuo et al., Identification of C2CD4A as a human diabetes susceptibility gene with a role in beta cell insulin secretion. Proc Natl Acad Sci U S A 116, 20033–20042 (2019).

13. L. Qiang, A. S. Banks, D. Accili, Uncoupling of acetylation from phosphorylation regulates FoxO1 function independent of its subcellular localization. J Biol Chem 285, 27396–27401 (2010).

14. A. S. Banks et al., Dissociation of the glucose and lipid regulatory functions of FoxO1 by targeted knockin of acetylation-defective alleles in mice. Cell Metab 14, 587–597 (2011).

15. J. Y. Kim-Muller et al., FoxO1 Deacetylation Decreases Fatty Acid Oxidation in beta-Cells and Sustains Insulin Secretion in Diabetes. J Biol Chem 291, 10162–10172 (2016).

16. A. Nitzsche et al., RAD21 cooperates with pluripotency transcription factors in the maintenance of embryonic stem cell identity. PLoS One 6, e19470 (2011).

17. N. D. Heintzman et al., Distinct and predictive chromatin signatures of transcriptional promoters and enhancers in the human genome. Nat Genet 39, 311–318 (2007).

18. D. Accili, K. C. Arden, FoxOs at the crossroads of cellular metabolism, differentiation, and transformation. Cell 117, 421–426 (2004).

19. V. Haberle, A. Stark, Eukaryotic core promoters and the functional basis of transcription initiation. Nat Rev Mol Cell Biol 19, 621–637 (2018).

20. E. Calo, J. Wysocka, Modification of enhancer chromatin: what, how, and why? Mol Cell 49, 825–837 (2013).

21. R. Raisner et al., Enhancer Activity Requires CBP/P300 Bromodomain-Dependent Histone H3K27 Acetylation. Cell Rep 24, 1722–1729 (2018).

22. C. Y. McLean et al., GREAT improves functional interpretation of cis-regulatory regions. Nature Biotechnology 28, 495–501 (2010).

23. S. X. Lee et al., FoxO transcription factors are required for hepatic HDL cholesterol clearance. J Clin Invest 128, 1615–1626 (2018).

24. K. C. Ehrlich, M. Lacey, M. Ehrlich, Tissue-specific epigenetics of atherosclerosis-related ANGPT and ANGPTL genes. Epigenomics 11, 169–186 (2019).

25. C. M. Allan, S. Taylor, J. M. Taylor, Two hepatic enhancers, HCR.1 and HCR.2, coordinate the liver expression of the entire human apolipoprotein E/C-I/C-IV/C-II gene cluster. J Biol Chem 272, 29113–29119 (1997).

26. M. Matsumoto, A. Pocai, L. Rossetti, R. A. Depinho, D. Accili, Impaired regulation of hepatic glucose production in mice lacking the forkhead transcription factor Foxo1 in liver. Cell Metab 6, 208–216 (2007).

27. R. A. Haeusler, K. H. Kaestner, D. Accili, FoxOs function synergistically to promote glucose production. J Biol Chem 285, 35245–35248 (2010).

28. R. A. Haeusler et al., Integrated control of hepatic lipogenesis versus glucose production requires FoxO transcription factors. Nature communications 5, 5190 (2014).

29. I. Goldstein et al., Transcription factor assisted loading and enhancer dynamics dictate the hepatic fasting response. Genome Res 27, 427–439 (2017).

30. M. Boergesen et al., Genome-wide profiling of liver X receptor, retinoid X receptor, and peroxisome proliferator-activated receptor alpha in mouse liver reveals extensive sharing of binding sites. Mol Cell Biol 32, 852–867 (2012).

31. S. Patel et al., GDF15 Provides an Endocrine Signal of Nutritional Stress in Mice and Humans. Cell Metab 29, 707–718 e708 (2019).

32. Y. Takeuchi et al., KLF15 Enables Rapid Switching between Lipogenesis and Gluconeogenesis during Fasting. Cell Rep 16, 2373–2386 (2016).

33. A. Vikram, G. Jena, S961, an insulin receptor antagonist causes hyperinsulinemia, insulin-resistance and depletion of energy stores in rats. Biochem Biophys Res Commun 398, 260–265 (2010).

34. H. V. Lin, D. Accili, Hormonal regulation of hepatic glucose production in health and disease. Cell Metab 14, 9–19 (2011).

35. M. S. Brown, J. L. Goldstein, Selective versus total insulin resistance: a pathogenic paradox. Cell Metab 7, 95–96 (2008).

36. S. Baldi et al., Influence of apolipoproteins on the association between lipids and insulin sensitivity: a cross-sectional analysis of the RISC Study. Diabetes Care 36, 4125–4131 (2013).

37. P. M. Titchenell, Q. Chu, B. R. Monks, M. J. Birnbaum, Hepatic insulin signalling is dispensable for suppression of glucose output by insulin in vivo. Nature communications 6, 7078 (2015).

38. V. T. Samuel et al., Fasting hyperglycemia is not associated with increased expression of PEPCK or G6Pc in patients with Type 2 Diabetes. Proc Natl Acad Sci U S A 106, 12121–12126 (2009).

39. A. Kalvisa et al., Insulin signaling and reduced glucocorticoid receptor activity attenuate postprandial gene expression in liver. PLOS Biology 16, e2006249 (2018).

40. R. Andersson, A. Sandelin, Determinants of enhancer and promoter activities of regulatory elements. Nat Rev Genet 10.1038/s41576-019-0173-8 (2019).

41. M. A. Zabidi et al., Enhancer-core-promoter specificity separates developmental and housekeeping gene regulation. Nature 518, 556–559 (2015).

42. M. M. Brent, R. Anand, R. Marmorstein, Structural basis for DNA recognition by FoxO1 and its regulation by posttranslational modification. Structure 16, 1407–1416 (2008).

43. X. C. Dong et al., Inactivation of hepatic Foxo1 by insulin signaling is required for adaptive nutrient homeostasis and endocrine growth regulation. Cell Metab 8, 65–76 (2008).

44. N. Kubota et al., Dynamic functional relay between insulin receptor substrate 1 and 2 in hepatic insulin signaling during fasting and feeding. Cell Metab 8, 49–64 (2008).

45. M. Lu et al., Insulin regulates liver metabolism in vivo in the absence of hepatic Akt and Foxo1. Nat Med 18, 388–395 (2012).

46. H. Yoshida et al., The cis-Regulatory Atlas of the Mouse Immune System. Cell 176, 897–912 e820 (2019).

47. M. Pawlak, P. Lefebvre, B. Staels, Molecular mechanism of PPARα action and its impact on lipid metabolism, inflammation and fibrosis in non-alcoholic fatty liver disease. Journal of Hepatology 62, 720–733 (2015).

48. A. Montagner et al., Liver PPARalpha is crucial for whole-body fatty acid homeostasis and is protective against NAFLD. Gut 65, 1202–1214 (2016).

49. T. Kuo et al., Induction of alpha cell-restricted Gc in dedifferentiating beta cells contributes to stress-induced beta-cell dysfunction. JCI Insight 5 (2019).

50. C. Postic et al., Dual Roles for Glucokinase in Glucose Homeostasis as Determined by Liver and Pancreatic β Cell-specific Gene Knock-outs Using Cre Recombinase. Journal of Biological Chemistry 274, 305–315 (1999).

51. J. R. Cook, F. Langlet, Y. Kido, D. Accili, Pathogenesis of Selective Insulin Resistance in Isolated Hepatocytes. Journal of Biological Chemistry 290, 13972–13980 (2015).

52. B. Langmead, S. L. Salzberg, Fast gapped-read alignment with Bowtie 2. Nat Methods 9, 357–359 (2012).

53. J. Feng, T. Liu, B. Qin, Y. Zhang, X. S. Liu, Identifying ChIP-seq enrichment using MACS. Nat Protoc 7, 1728–1740 (2012).

54. S. Heinz et al., Simple combinations of lineage-determining transcription factors prime cis-regulatory elements required for macrophage and B cell identities. Mol Cell 38, 576–589 (2010).

55. A. R. Quinlan, BEDTools: The Swiss-Army Tool for Genome Feature Analysis. Curr Protoc Bioinformatics 47, 11 12 11–34 (2014).

56. E. P. Consortium, An integrated encyclopedia of DNA elements in the human genome. Nature 489, 57–74 (2012).

