## Supplemental files for "An integrative transcriptional logic model of hepatic insulin resistance"

**SUPPLEMENTAL INFORMATION**

This appendix has been provided by the authors to give additional information about the present study.

**Figures**

Figure S1. In vivo kinetics of hepatic FoxO1 localization related to Figure 1

Figure S2. Optimization of FoxO1 ChIP-seq related to Figure 1

Figure S3. Basic characteristics of FoxO1 binding sites related to Figure 1

Figure S4. The top 10 motifs found from the HOMER *de novo* motif analysis related to Figure 1

Figure S5. Comparison of FoxO1 active enhancers and enhancer-less promoter/TSS in fast and refeed related to Figure 2

Figure S6. Linear regression analysis of FoxO1 intron peaks related to Figure 3

Figure S7. Experimental diagram for resilience analysis of FoxO1-regulated genes related to Figure 4

Figure S8. Representative images of FoxO1 peaks in insulin resistance related to Figure 6

**Tables**

Table S1. Peaks associated with significant gene expression related to Figure 1f

Table S2. FoxO1 peaks in introns related to Figure 3

Table S3. DEGs in A-FLKO and C-FLKO related to Figure 4

Table S4. The correlation of FoxO1 binding with liver expressed genes related to Figure 5

Table S5. Active enhancer FoxO1 and PPARα peaks associated with DEGs in A-FLKO related to Figure 5

Table S6. Technical information for FoxO1-GFP ChIP-seq related to Method

### SUPPLEMENTAL FIGURE LEGENDS

Figure S1. In vivo kinetics of hepatic FoxO1 localization

(a–b) Time- (a) and dose- (b) dependence of FoxO1 nucleocytoplasmic translocation in response to exogenous insulin. The images show liver FoxO1and HNF4α immunohistochemistry before and 3, 15, 30, and 60 min after intra-portal insulin injection (0.1U/kg) (a), and 15 min after injection of 0.1U/kg, 0.01 U/kg, and 0.001 U/kg, respectively (b). Scale bar = 20 μm. (c) FoxO1 and HNF4α immunohistochemistry in liver during 16-hr-fasted/refed time course. (d) Changes to plasma glucose and insulin levels during 4-hr- or 16-hr-fasting, and followed by refeeding (n = 5 mice per data point). Data are shown as mean ± SD. Squares: glucose levels (mg/dl); Circles: plasma insulin levels (ng/μl); Solid arrows: timing of refeeding; Red filled squares and green filled circles: post-prandial time points (e) Western blot of phospho-Akt, pan-Akt and β-actin in the same time points shown in (d).

Figure S2. Optimization of FoxO1 ChIP-seq and basic characteristics of FoxO1 binding sites

(a) Western blot of FoxO1 and β-actin in primary hepatocyte and liver of wild type control (WT) or FoxO1-Venus (FxV) mice. (b) ChIP-Western blot of GFP antibody or IgG in liver of FoxO1-Venus mice. (c) ChIP-qPCR with FoxO1 or GFP antibody for known FoxO1 target genes. (d–h) IGV genome browser view of FoxO1 ChIP-seq for representative genes, such as Igfbp1 (d), G6PC (e), Pck1 (f), and Fkbp5(g), and corresponding results of ChIP-qPCR with GFP antibody in fasted and refed condition (h). red bar showed the sequence amplified by the primers in ChIP-qPCR.

Figure S3.

(a–b) Circo plots (a) and TSS-center histogram (b) of FoxO1 peaks in fasted or refed conditions from Figure 1. (c) Known motif analysis of FoxO1 ChIP-seq. Logos of the recovered FoxO1 motif shown as in Figure 1e (p = 1e-156 in fast, 1e-166 in refeed). (d–e) Frequency of FoxO1 motifs relative to FoxO1 peaks in fast or refeed (d) and TSS (e). (f) mRNA measured by qPCR of liver of AAV-GFP- or AAV-Cre-injected mice (n = 5 mice per group) Data are shown as mean ± SEM. (g) Western blot of FoxO1 and β-actin in liver, quadriceps muscle (Quad), and epididymal fat (eWAT) in AAV-GFP- or AAV-Cre-injected mice.

Figure S4. The top 10 motifs found from the HOMER *de novo* motif analysis

The results of motif analysis through HOMER *de novo* motif finding program in fasted and refed condition.

Figure S5. Comparison of FoxO1 active enhancers and enhancer-less promoter/TSS in fast and refeed

(a) Normalized mean signal intensity for each ChIP-seq read: FoxO1 in fast (cyan line) or refeed (red line), H3K27ac in fast (red dotted line) or refeed (blue dot line), and H3K4me1 (green dotted line) centered on TSS. (b) Venn diagram of the number of FoxO1 active enhancer peaks in fasted or refed conditions. (c) List of GO in blue circles in Figure 2b and their -log10 FDR. (d) Linear regression analysis between FoxO1 active enhancers in fast vs. refeed. (e, f) Same GO and linear regression analysis applied to enhancer-less promoter/TSS. (g) Linear regression analysis between H3K27ac peaks between fast vs. refeed.

Figure S6. Linear regression analysis of FoxO1 intron peaks

Linear regression analysis of FoxO1 enhancer-less (a) or active enhancers (b) introns between fast vs. refeed.

Figure S7. Experimental diagram for resilience analysis of FoxO1-regulated genes

For adult-onset liver-specific FoxO1 knockout mice (A-FLKO), we injected AAV8.TBG.eGFP or AAV8 TBG.Cre in 5–7 weeks FoxO1lox/lox mice and took liver from 4-hr-fasted or 4hr-fasted/1-hr-refed mice 3 weeks after injection. For constitutive FoxO1 knockout mice (C-FLKO), we took liver tissues from 8–10 weeks FoxO1lox/lox or Alb-Cre; FoxO1lox/lox mice.

Figure S8. Representative images of FoxO1 peaks in insulin resistance

IGV Genome browser views of FoxO1 peaks with or without S961 treatment and associated H3K27ac and H3K4me1 marks of at KLF15 (a), ApoC4/C2/C1/E (b), and Scarb1 (c).

Table S4. The correlation of FoxO1 binding with liver expressed genes

| | FoxO1 unique peaks | FoxO1 peaks co-binding with the other TFs | *p* Value |
|---|---|---|---|
| Active Enhancer (number of peaks) | 1370 | 2833 | |
| Peaks regulating gene expression [peak number, (%)] | 153 (11.2 %) | 153 (6.1 %) | |
| Peaks without gene expression changes [peak number, (%)] | 1217 (88.8 %) | 2680 (94.6 %) | < 0.0001 |
| Non Active Enhancer (number of peaks) | 4268 | 5347 | |
| Peaks regulating gene expression [peak number, (%)] | 260 (6.1 %) | 317 (5.9 %) | |
| Peaks without gene expression changes [peak number, (%)] | 4008 (93.9 %) | 5030 (94.1 %) | 0.7623 |

Combined analysis of FoxO1 binding sites with or without active enhancer marks and gene expression profiles near FoxO1 binding sites. The data from Fasted condition was used for this analysis, and the genes showing FDR < 0.1 between A-FLKO and WT control were assigned as FoxO1 regulating genes. FoxO1 peaks were assigned into two groups of FoxO1 unique binding sites or the sites bound by FoxO1 with any of the other TFs of CREB, GR, or PPARα. Chi-square test was used for statistical analysis.

Table S6. . Technical information for FoxO1-GFP ChIP-seq related to Method

| | FoxO1-GFP_Vehicle | | FoxO1-GFP_S961 | | H3K27ac_Vehicle | |
|---|---|---|---|---|---|---|
| | Fast | Refeed | Fast | Refeed | Fast | Refeed |
| ChIP-seq replicates | Biological triplicate | Biological triplicate | Biological triplicate | Biological triplicate | Biological triplicate | Biological triplicate |
| Total number of reads | 39,082,642 | 37,790,703 | 36,916,846 | 41,400,137 | 35664103 | 32759403 |
| Number of uniquely aligned reads | 30,977,910 | 31,391,870 | 29,241,530 | 34,681,135 | 29838325 | 27403118 |
| Percentage of aligned reads (%) | 89.65 | 94.93 | 88.51 | 93.45 | 92.42 | 92.57 |

Supplemental Figure 1

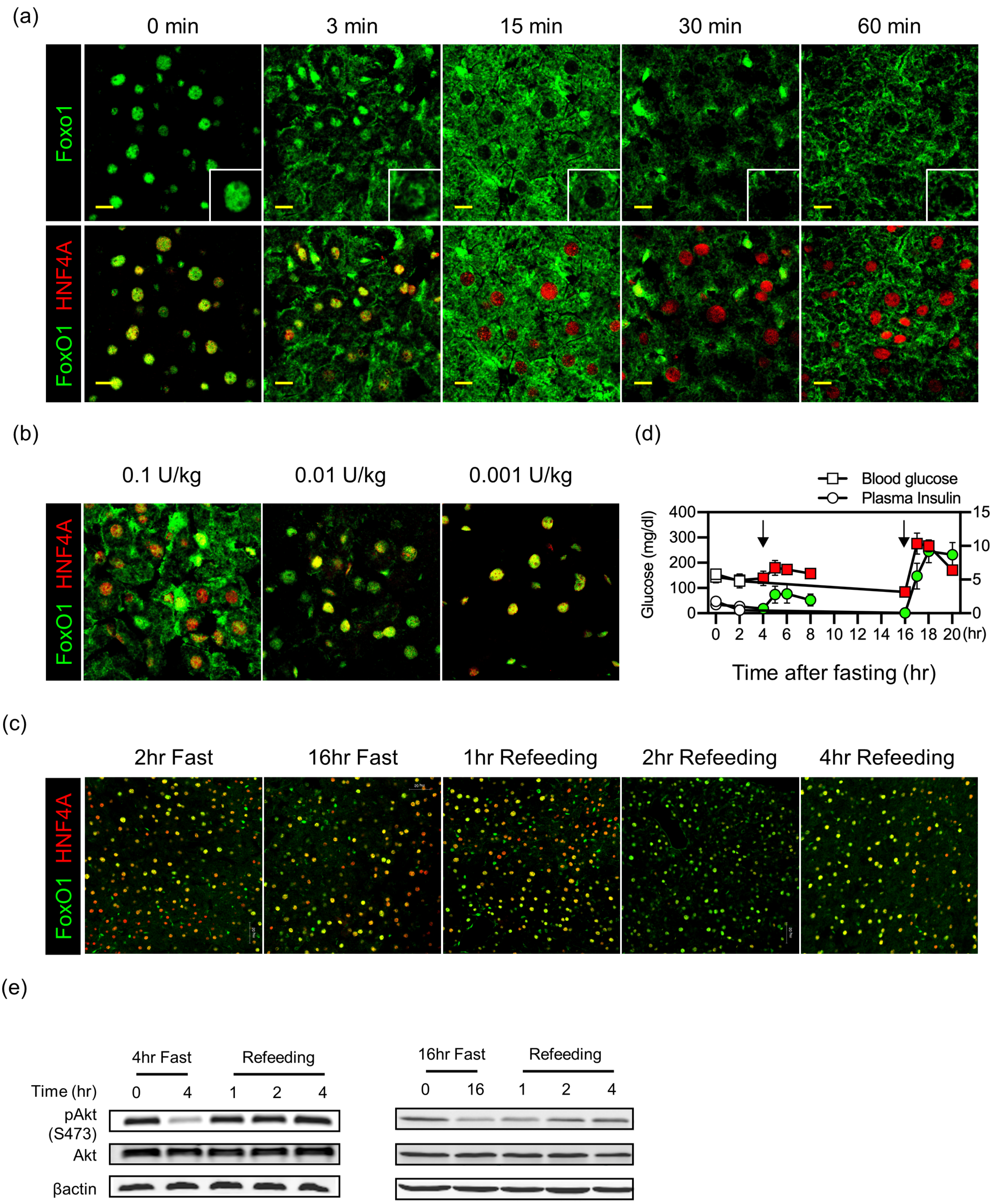

Supplemental Figure 2

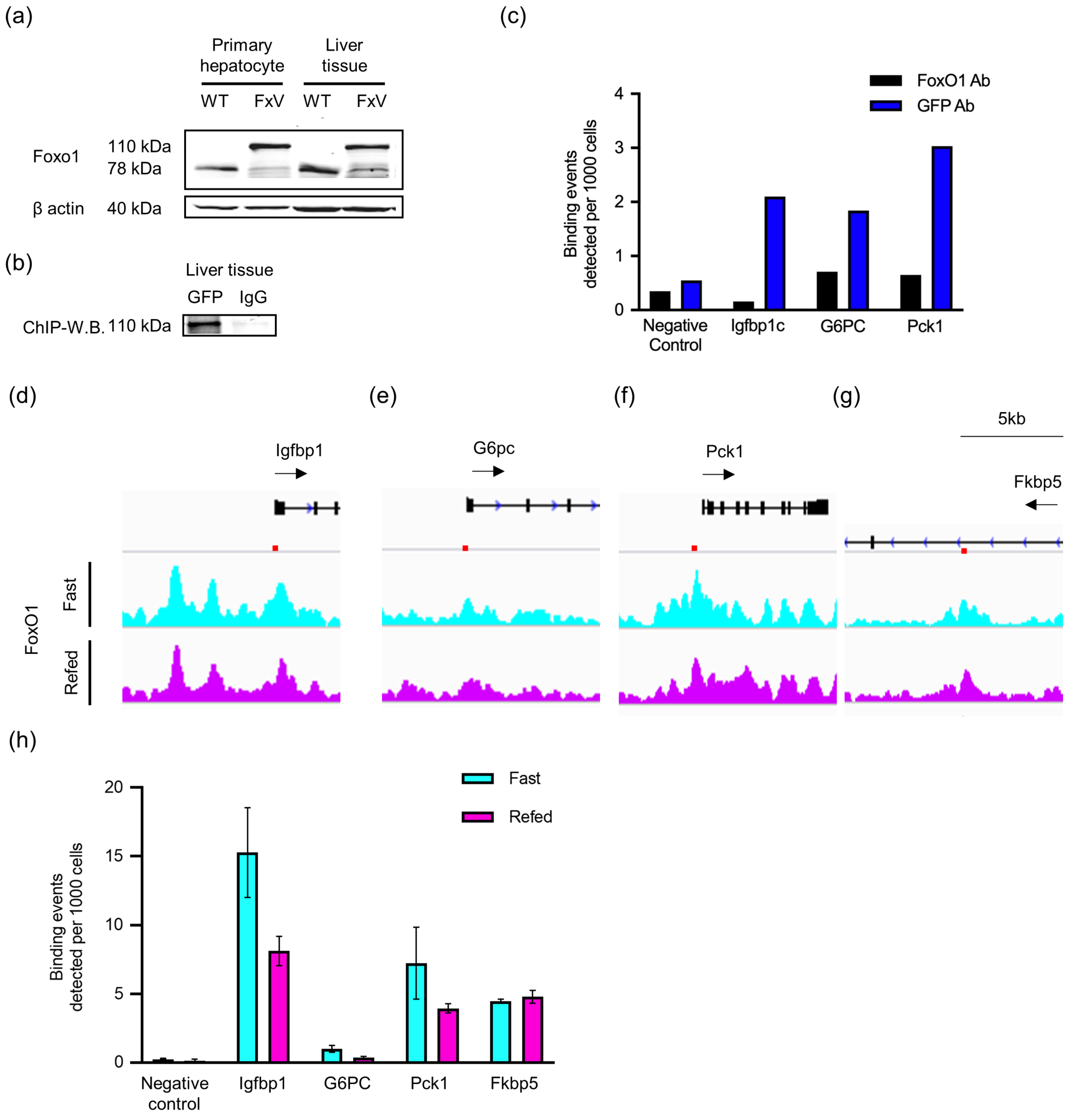

Supplemental Figure 3

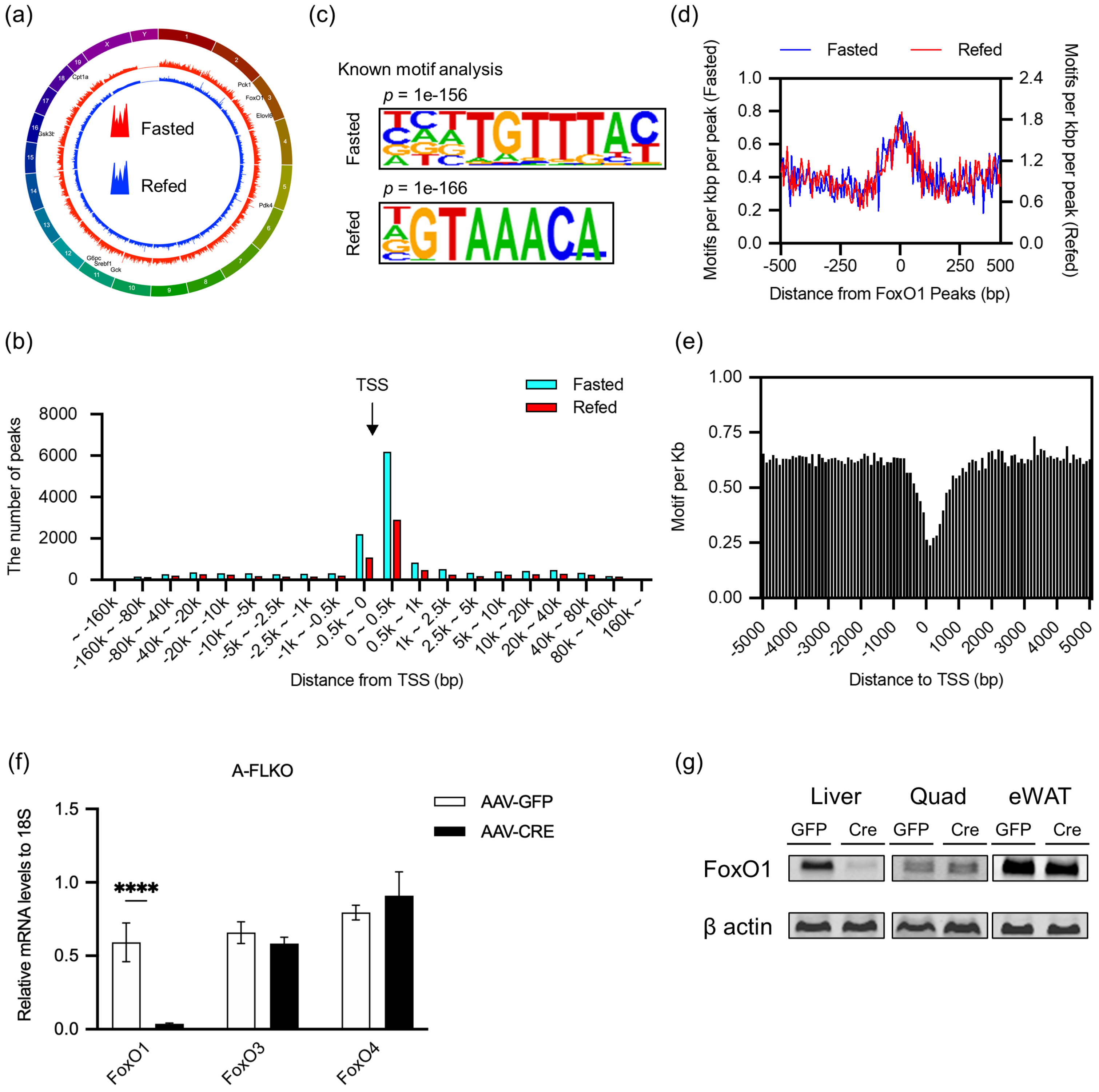

Supplemental Figure 4

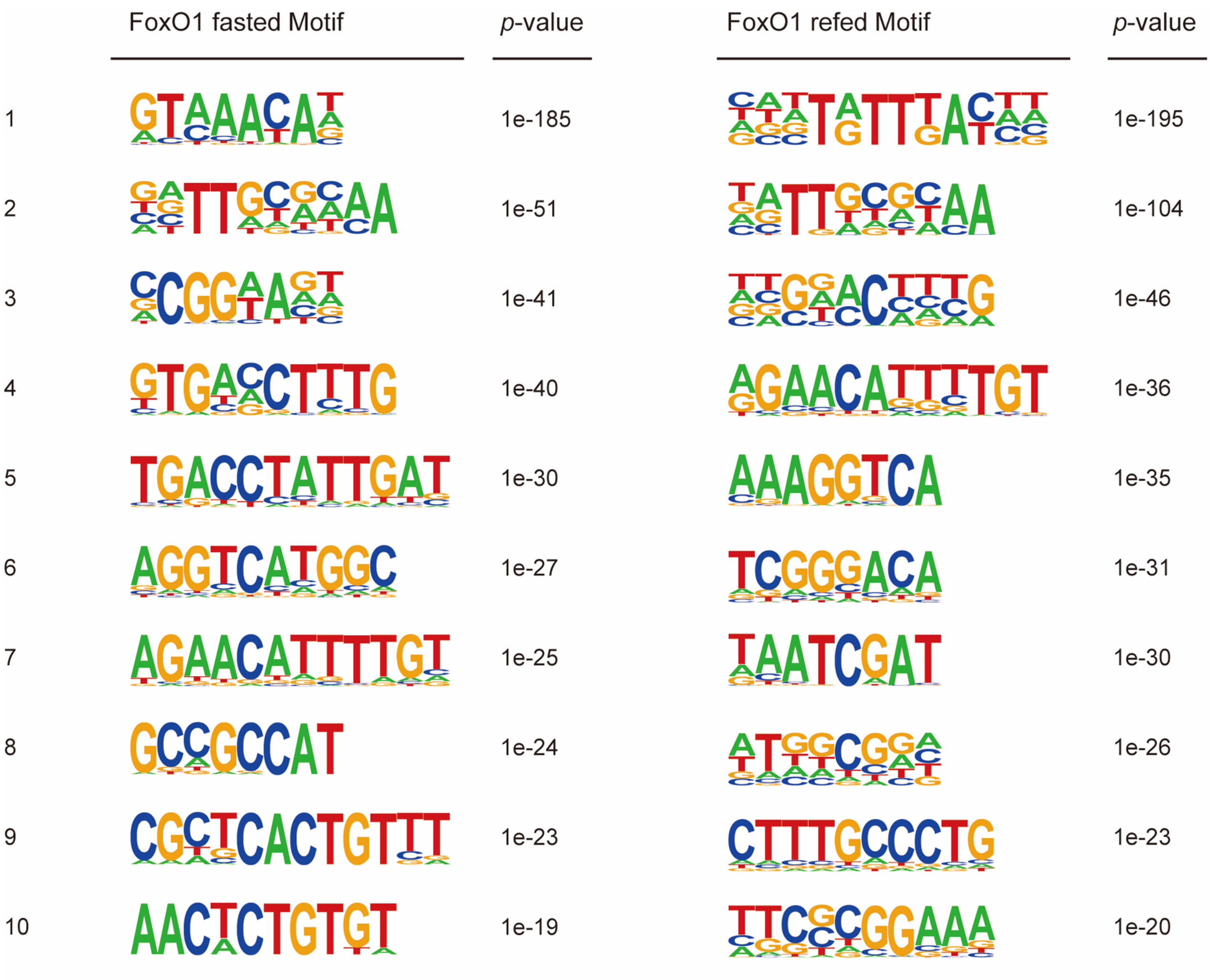

Supplemental Figure 5

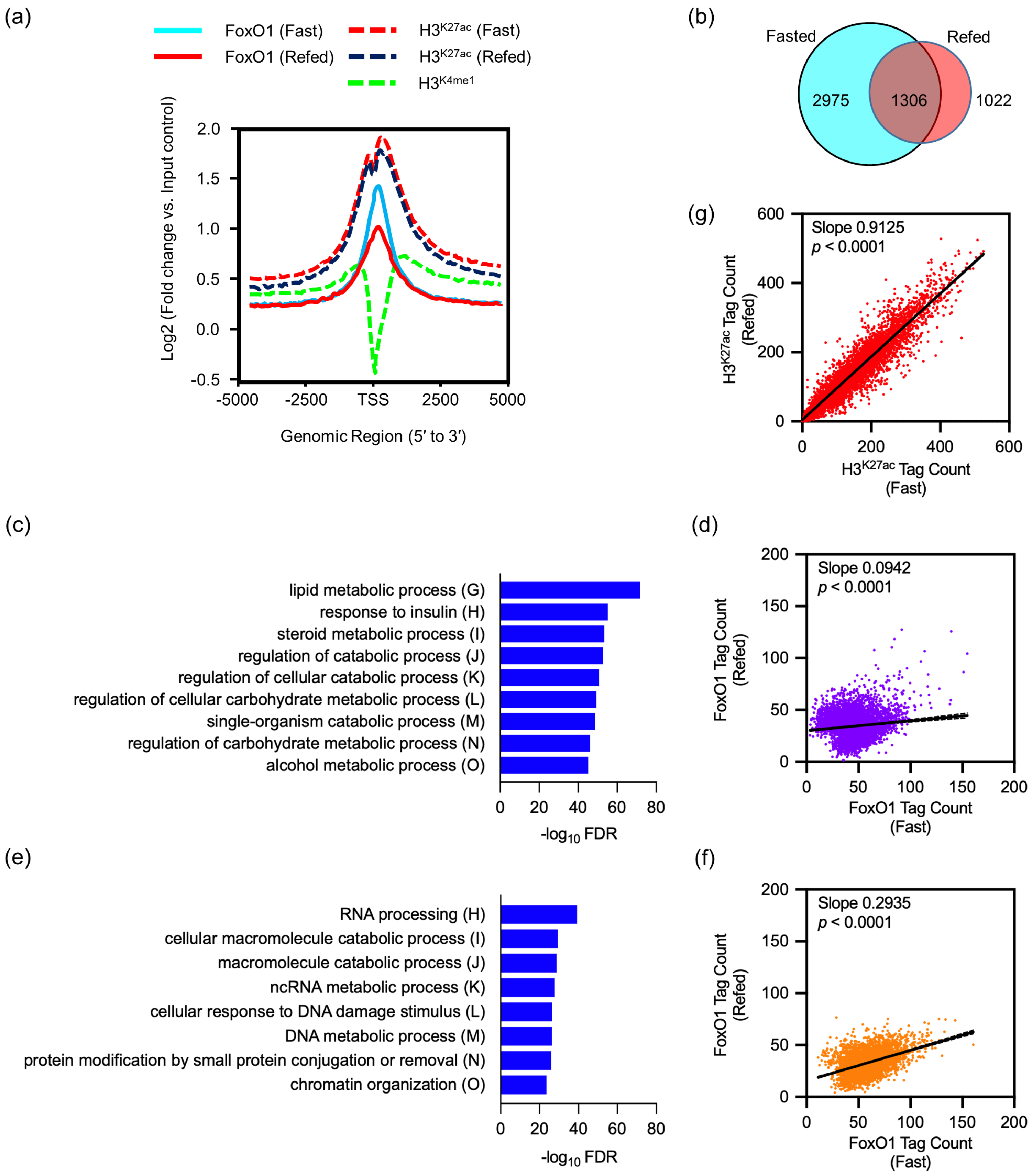

Supplemental Figure 6

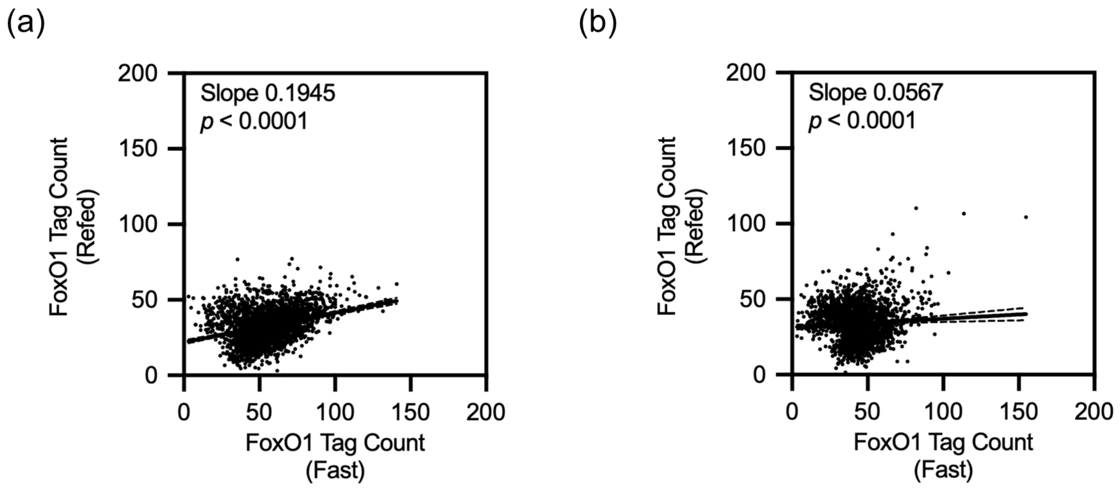

Supplemental Figure 7

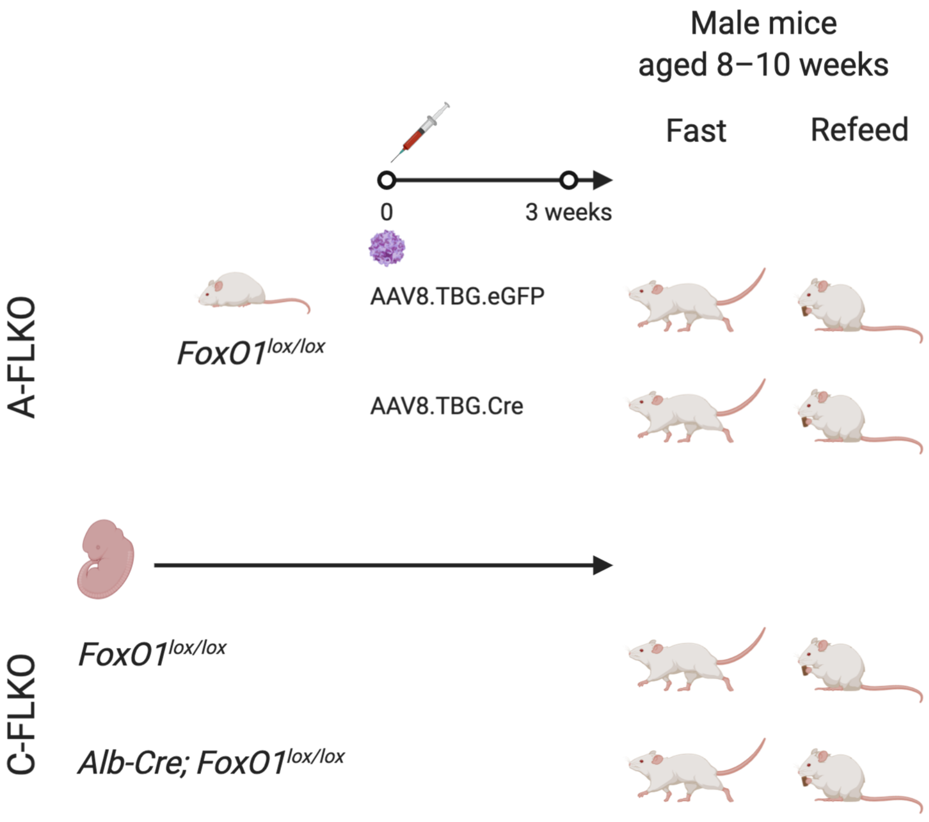

Supplemental Figure 8

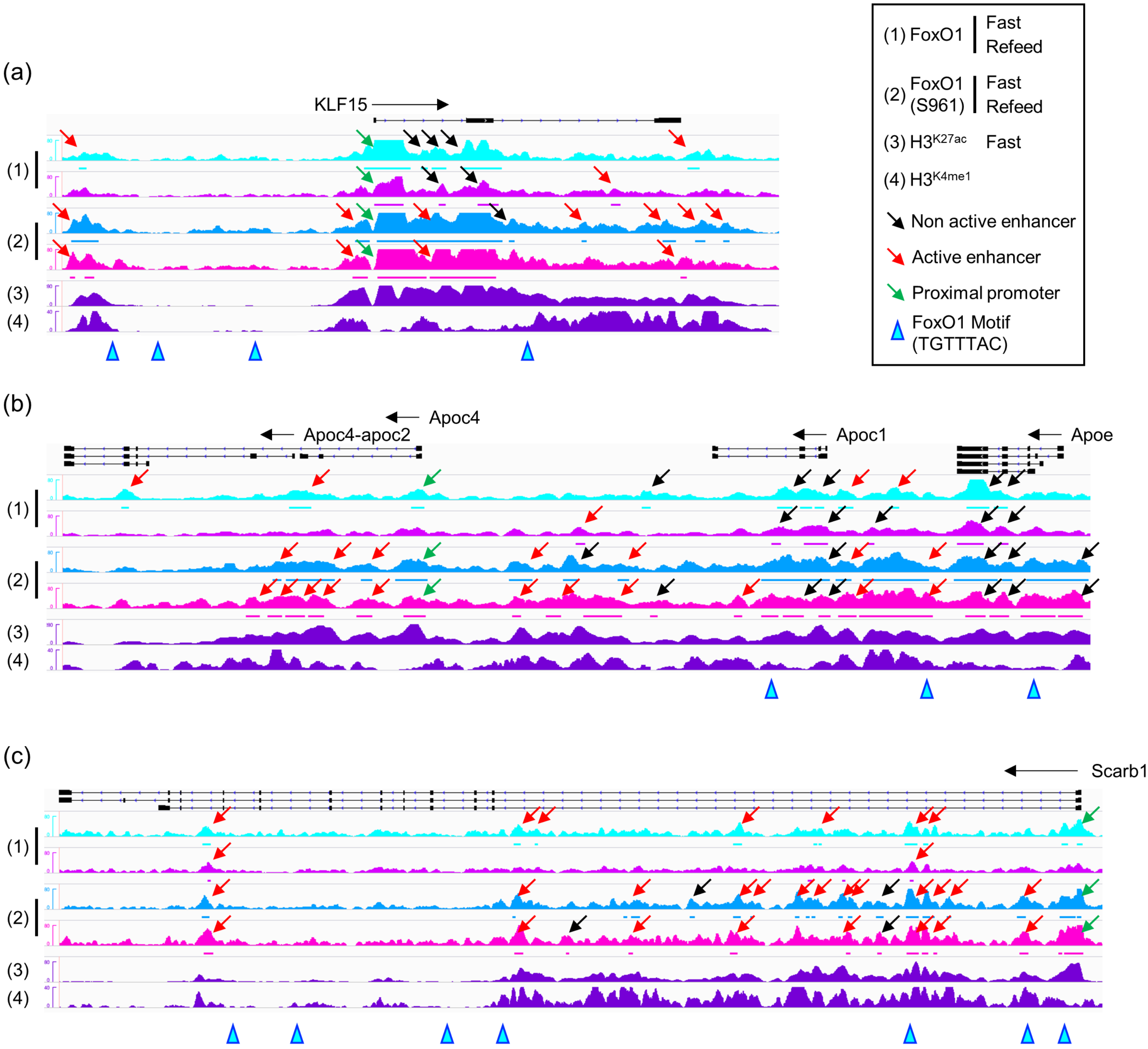
